# Identification of Fis1 interactors in *Toxoplasma gondii* reveals a novel protein required for peripheral distribution of the mitochondrion

**DOI:** 10.1101/805911

**Authors:** Kylie Jacobs, Robert Charvat, Gustavo Arrizabalaga

## Abstract

*Toxoplasma gondii*’s singular mitochondrion is very dynamic and undergoes morphological changes throughout the parasite’s life cycle. During parasite division, the mitochondrion elongates, enters the daughter cells just prior to cytokinesis and undergoes fission. Extensive morphological changes also occur as the parasite transitions from the intracellular to the extracellular environment. We show that treatment with the ionophore monensin causes reversible constriction of the mitochondrial outer membrane, and that this effect depends on the function of the fission related protein Fis1. We also observed that mislocalization of the endogenous Fis1 causes a dominant negative effect that affects the morphology of the mitochondrion. As this suggests Fis1 interacts with proteins critical for maintenance of mitochondrial structure, we performed various protein interaction trap screens. In this manner we identified a novel outer mitochondrial membrane protein, LMF1, which is essential for positioning of the mitochondrion in intracellular parasites. Normally, while inside a host cell, the parasite mitochondrion is maintained in a lasso shape that stretches around the parasite periphery where it has regions of coupling with the parasite pellicle, suggesting the presence of membrane contact sites. In intracellular parasites lacking LMF1 the mitochondrion is retracted away from the pellicle and instead is collapsed, as only normally seen in extracellular parasites. We show that this phenotype is associated with defects in parasite fitness and mitochondrial segregation. Thus, LMF1 is necessary for mitochondrial association with the parasite pellicle during intracellular growth and proper mitochondrial morphology is a prerequisite for mitochondrial division.

**IMPORTANCE:** *Toxoplasma gondii* is an opportunistic pathogen that can cause devastating tissue damage in the immunocompromised and the congenitally infected. Current therapies are not effective against all life stages of the parasite and many cause toxic effects. The single mitochondrion of this parasite is a validated drug target and it changes its shape throughout its life cycle. When the parasite is inside of a cell, the mitochondrion adopts a lasso shape that lies in close proximity to the pellicle. The functional significance of this morphology is not understood nor are the proteins involved currently known. We have identified a protein that is required for proper mitochondrial positioning at the periphery and that likely plays a role in tethering this organelle. Loss of this protein results in dramatic changes to the mitochondrial morphology and significant parasite division and propagation defects. Our results give important insight into the molecular mechanisms regulating mitochondrial morphology.

## INTRODUCTION

*Toxoplasma gondii* is an opportunistic protozoan parasite that can infect nearly any nucleated cell in a wide range of warm-blooded organisms. This promiscuity contributes to *Toxoplasma gondii* being one of the most widespread and successful parasites in the world. It has been estimated that approximately one-third of the world’s human population is infected with *Toxoplasma* (1)*. Toxoplasma* infections in humans are usually the result of the ingestion of parasite oocysts deposited in the feces of felines or of tissue cysts present in raw or undercooked meat from infected warm-blooded animals. Despite all of these routes of infection, there are generally few side effects associated with toxoplasmosis because its acute state is susceptible to the healthy immune system. However, the parasite can evade the immune response by converting to a latent encysted form, thus establishing a chronic infection. In immunocompromised individuals and lymphoma patients, new infections or re-activation of pre-existing cysts can lead to toxoplasmic encephalitis, among other complications. Additionally, in congenital infections, toxoplasmosis can lead to blindness, severe neurological problems or even death given the immature nature of the fetal immune system. Although there are drugs that target acute toxoplasmosis, they are often toxic and all are ineffective against the chronic stage. Therefore, it is important to identify novel and unique drug targets that are effective against multiple stages of the parasite life cycle.

One interesting feature of *Toxoplasma* is its singular mitochondrion, which is very large and extends to the periphery of the cell. In addition to its plant-like features, such as tubular cristae, the *Toxoplasma* mitochondrion has a streamlined mitochondrial genome only encoding 3 proteins: c*ox1, cob*, and *cox3* (2). These unique features of the mitochondrion, along with its essentiality for parasite survival, make it an interesting drug target. The clinical effectiveness of the mitochondrial inhibitor atovaquone against *Toxoplasma* and related parasites, highlights the validity of this organelle as a target for anti-parasitic therapy (3). Atovaquone acts by mimicking ubiquinone and competitively binding to the cytochrome bc_1_ complex, which blocks the electron transport chain and prevents energy production in the mitochondrion (4). Other electron transport chain inhibitors, such as the endochin-like quinolones, are still being studied for efficacy in *Toxoplasma* and *Plasmodium* (5, 6). Aspects of the apicomplexan mitochondrion that remains unexplored as a potential target are its morphology and division. Chemical inhibition of mitochondrial fission has been shown to have cytoprotective effects in cardiovascular injury models (7) and to prevent cell proliferation in lung cancer (8) and glioblastoma (9). Thus, in depth understanding of the regulation of mitochondrial morphology and dynamics in *Toxoplasma*, could reveal novel therapeutic targets.

The mitochondrion of *Toxoplasma* is highly dynamic and exhibits significant morphological changes during the parasite’s lifecycle and in response to various stressors. As there is only one mitochondrion per parasite, its division is tightly coordinated with the division of the rest of the parasite (10). *Toxoplasma* divides by endodyogeny, a specialized process through which two daughter cells form within the mother parasite and during which each organelle is either made de novo or elongated and divided for incorporation into the daughter parasites. The mitochondrion divides very late in this process and is not incorporated into the daughter parasites until the parasites have almost completely emerged from the mother parasite (3). The proteins involved in the budding, division, and segregation of mitochondrial material have not been thoroughly examined and *Toxoplasma gondii* is lacking almost all of the homologues of proteins used in these processes. For example, the mitochondrial division machinery is made up of three components: a fission protein to recruit proteins necessary for division, adaptor proteins to provide a scaffold, and a dynamin-related protein to cause the final scission of the mitochondrion (11). *Toxoplasma* encodes one homolog for the fission protein, Fis1 (TGGT1_263323) and three potential dynamin related proteins (Drps): DrpA, DrpB, and DrpC. TgDrpA and TgDrpB have been shown to be required for apicoplast replication and secretory organelle biogenesis, respectively (12, 13). TgDrpC is divergent from the typical Drp due to the absence of a conserved GTPase Effector Domain, which is generally required for function. We recently showed that TgDrpC interacts with proteins that exhibit homology to those involved in vesicle transport (14, 15). Additionally, TgDrpC localizes to cytoplasmic puncta that redistribute to the growing edge of the daughter parasites during endodyogeny. Loss of TgDrpC stalls the division process and leads to rapid deterioration of multiple organelles, including the mitochondrion. Independent work also shows that this loss halts mitochondrial division (16). Therefore, TgDrpC appears to contribute to multiple processes including vesicular trafficking, organelle stability, division, and potentially mitochondrial division.

After repeated division cycles, the parasites egress from the cell and are exposed to the extracellular environment, where the mitochondrion alters its morphology. When *Toxoplasma* is within a host cell it maintains its mitochondrion in a lasso shape that spans the parasite’s periphery and is adjacent to the parasite pellicle (2, 10, 17). Immediately after egress the mitochondrion retracts from the periphery of the parasite and transitions to a “sperm-like” morphology, where the majority of mitochondrial material is at the apical end of the parasite with a tail of material extending towards the basal end (17). Prolonged exposure to the extracellular environment, results in transition to a completely collapsed mitochondrion. Upon reinvasion, the mitochondrion returns to the “lasso” shape almost immediately (17). Electron microscopy of parasites with lasso shaped mitochondrion reveals the presence of regions of close abutment between the outer mitochondrion membrane (OMM) and the inner membrane complex (IMC), in which the membranes retain a constant distance over stretches of 100 nm– 1000 nm (17). The average distance between the OMM and the IMC was calculated to be approximately 25 nm, which would suggest the presence of membrane contact sites (18, 19). Neither the functional significance nor the components of the proposed contact between the mitochondrion and the pellicle are known.

We have also observed that the mitochondrion of *Toxoplasma* significantly changes its morphology in response to exposure to the anti-coccidial drug monensin. Monensin is a sodium hydrogen exchanger that induces oxidative stress (20) and autophagic cell death (21). We demonstrate that monensin’s effect on the mitochondrion morphology is reversible, suggesting that *Toxoplasma* has mechanisms to rearrange the mitochondrial structure in response to drug induced stress. As the mitochondrion appears broken down upon monensin treatment we investigated the role of the fission machinery on this phenomenon. Here, we show that monensin induces a reversible constriction of the outer mitochondrial membrane and that this effect is in part dependent on the fission protein Fis1. We also show that, although Fis1 is not required for parasite survival, mislocalization of Fis1 away from the outer mitochondrial membrane results in aberrant mitochondrial morphology. We hypothesize that the dominant negative effect caused by mislocalization of Fis1 is due to misdirecting critical proteins away from the mitochondrion. Accordingly, we identified interactors of Fis1. One such interactor, TgGT1_265180, proved to be required for parasite growth, division, and mitochondrial segregation. Importantly, the mitochondria of intracellular parasites lacking this Fis1 interactor are not lasso shaped, but instead are collapsed away from the parasite periphery. Accordingly, we hypothesize that this novel protein is part of the proposed scaffold that mediates membrane contact sites between the mitochondrion and the parasite pellicle.

## RESULTS

### Monensin-induced mitochondrial remodeling is reversible

We had previously observed that treatment with the polyether ionophore monensin induced gross morphological changes in *Toxoplasma*, including alterations in the Golgi apparatus and mitochondrion (20). In particular, the mitochondrion, which under normal growth conditions appears as a lasso along the periphery of the parasite, becomes fragmented in appearance upon monensin treatment (Fig. 1A). To assess whether this effect on the parasite mitochondrion was reversible, parasites were treated with either vehicle or 1 mM monensin for 12 hours followed with a 12 hours recovery period on normal growth medium. Under vehicle-treated conditions, parasites exhibited intact mitochondria in greater than 91% of vacuoles (Figs. 1A and B). By contrast, following 12 hours of monensin treatment, only 6.25±11.8% of vacuoles contained parasites with intact mitochondria, congruent with previous findings (Figs. 1A and B). Interestingly, this phenotype is reversed when the drug is removed and parasites are allowed to recover for 12 hours. After the 12-hour recovery period, in 79.5±5.5% of vacuoles all parasites show normal mitochondrial morphology (Figs. 1A and B). Of note, there was no observed reduction in total number of parasite-containing vacuoles between the cultures for which the drug was removed and those for which it was not, indicating a genuine recovery and not an expansion of surviving parasites.

**Figure 1.**
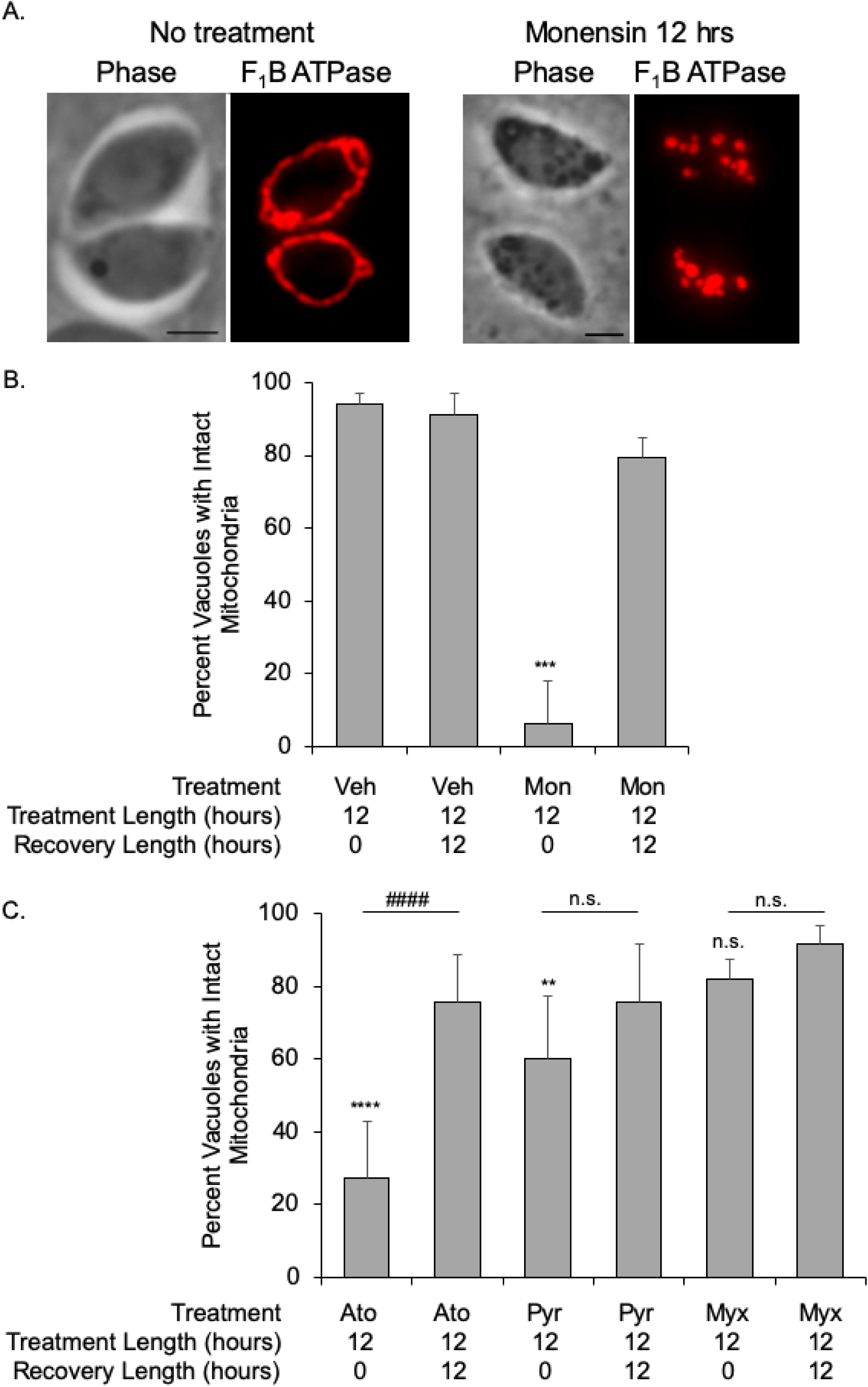
Drug-induced mitochondrial disruption is reversible. To determine the effect of drugs on mitochondrial morphology, intracellular parasites were treated with various agents. Parasite mitochondrial morphology was examined by visualizing the IMM localized F_1_B ATPase through immunofluorescence microscopy. A. Left panels show mitochondrion after treatment with vehicle, while right panels show effect of treatment with 1 mM monensin for 12 hours. Mitochondrion in the vehicle treated parasites shown is considered intact while in the drug treated parasites shown is considered disrupted. Scale bar, 2 µm. B. The percent of vacuoles with intact mitochondria is presented for parasites that were vehicle (veh) treated, monensin (mon) treated for 12 hours or monensin treated followed by a 12-hour recovery period. C. The effects of the anti-parasitic drug atovaquone (Ato, 100 nM), pyrimethamine (Pyr, 100 µM), and myxothiazol (Myx, 50 ng/mL) on the mitochondrion was assessed. With each drug we also tested the effect of a 12-hour recovery period after 12 hours of drug treatment. For all graphs 100 vacuoles for each condition were enumerated at random, and the data is presented as the average ± SD from 3 independent experiments. One-way ANOVA with post-hoc Tukey was utilized for statistical analysis. In B ****p<0.0001 in comparison to other treatments, in C each drug treatment was compared to vehicle, ****p<0.0001, **p<.007, and each treatment was compared to treatment followed by recovery, ###p<0.001.

To determine whether mitochondrial remodeling is a generalized drug response, parasites were challenged with atovaquone and myxothiazol, both cytochrome bc_1_ complex inhibitors, and pyrimethamine, a dihydrofolate reductase inhibitor not known to affect mitochondrial function. After 12 hours of atovaquone treatment, only 27.3±15.5% of parasites showed intact mitochondria and, just as what is observed with monensin, this effect was reversed by removal of drug and a 12-hour drug-free recovery period (75.8±13.0% intact mitochondria, Fig. 1C). By contrast, a lethal dose of myxothiazol (50 g/mL, (22, 23)) had little effect on mitochondrial morphology, with 82±5.4% of vacuoles with intact mitochondrion after treatment, which increased to 91.5±5.2% upon drug removal, although these effects did not meet statistical significance (Fig. 1C). Upon pyrimethamine treatment 60.3±17.0% of parasites had aberrant mitochondrial morphology. As this level did not change with statistical significance upon removal of drug (75.5±16.2%), the disruption of the mitochondrion observed is likely the consequence of parasite death and not the temporary and reversible rearrangement seen with monensin and atovaquone. Taken together, reversible mitochondrial disruption does not appear to be a generalized mechanism for responding to stress induced via drug challenge. Moreover, *Toxoplasma* possesses the capacity for reversible mitochondrial rearrangement in response to specific drug-induced stress.

### A fission protein homolog localizes to the outer mitochondrial membrane

The most striking aspect of monensin treatment in *Toxoplasma* is the disruption of mitochondrial morphology, producing what appears to be a fragmented organelle. A survey of the *Toxoplasma* database (ToxoDB) to identify homologs involved in mitochondrial dynamics revealed that the genome of *Toxoplasma* is rather bereft of proteins that participate in the fusion and fission processes. However, we were able to identify a protein (TGGT1_263323) with homology to the fission 1 (Fis1) protein from higher eukaryotes. TGGT1_263323, referred to hereafter as Fis1, is a 154 amino acids protein and contains two tetratricopeptide (TPR) domains, a C-terminal transmembrane (TM) domain followed by a three amino acid C-terminal sequence (CTS). In previous work focused on the characterization of membrane anchor domains in *Toxoplasma* we showed through transient transfection of an N-terminal HA tagged Fis1 that it localized to the mitochondrion (24). In order to further characterize the localization and function of Fis1, we established a parasite strain stably expressing an N-terminally HA epitope-tagged version of Fis1 (Fig. 2A). Immunofluorescence assay (IFA) of intracellular parasites of this strain (RHΔ*hpt*+HAFis1) confirmed that Fis1 localized to the parasite mitochondrion by co-localization with F_1_B ATPase protein, which is located in the inner mitochondrial membrane (IMM) (Fig. 2B). Super-resolution imaging shows that the Fis1 signal envelops the signal from F_1_B ATPase (Fig. 2C). This strongly suggests that, as expected for Fis1 proteins, Fis1 localizes to the outer mitochondrial membrane.

**Figure 2.**
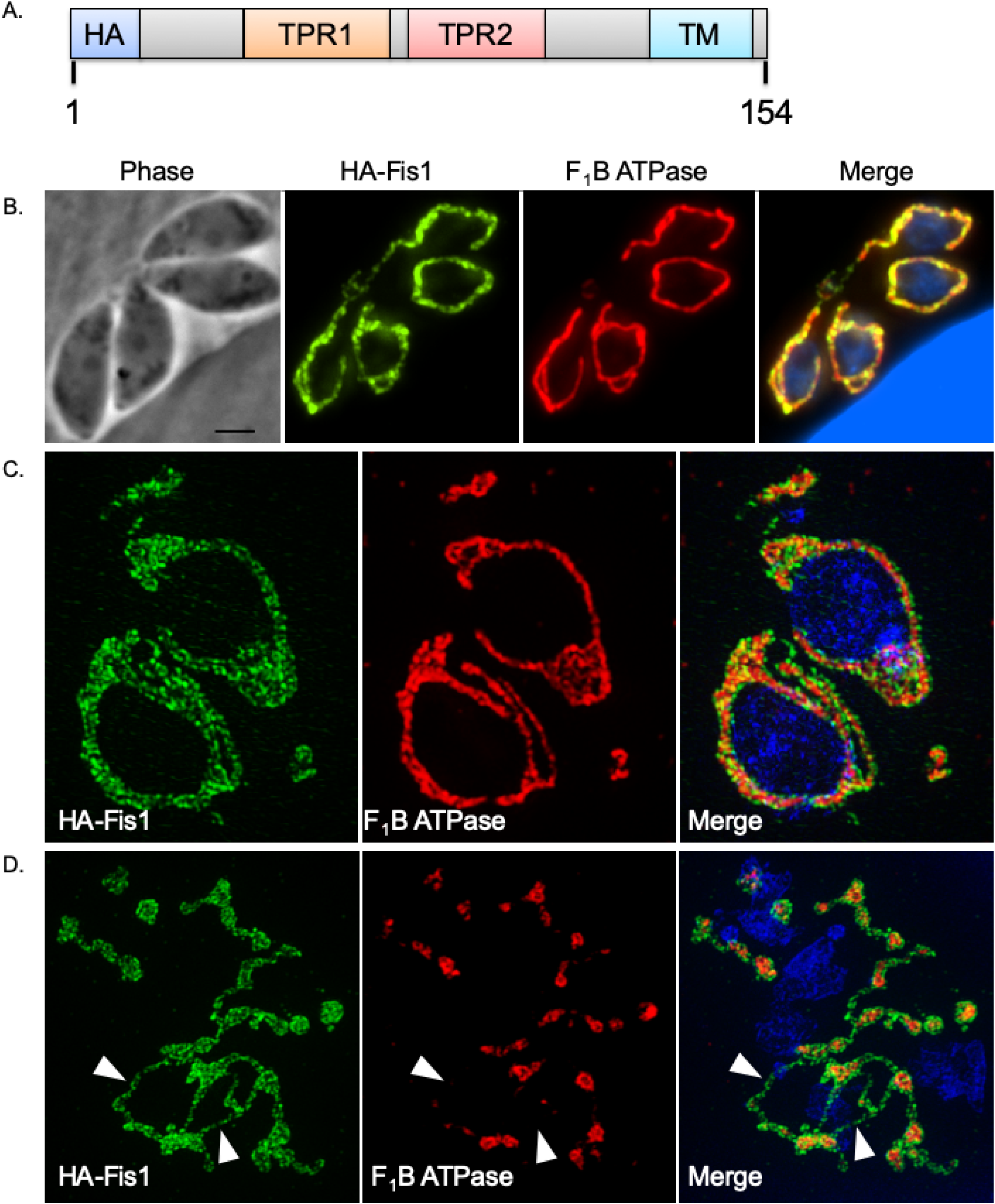
Fis1 localizes to the *Toxoplasma* outer mitochondrial membrane, which remains intact after monensin treatment. To determine the subcellular distribution of the fission protein homolog Fis1, a parasite strain expressing an ectopic copy of Fis1 including an N-terminal HA epitope tag was generated. A. Illustration shows the exogenously expressed epitope tagged Fis1. Protein domains in Fis1 are indicated: tetratricopeptide repeat domains (TPR) 1 and 2 and transmembrane (TM) domain. B-D. Intracellular parasites of the (HA)Fis1 expressing strain were analyzed by IFA using antibodies against the HA tag to detect Fis1 (in green) and against the *Toxoplasma* F_1_B ATPase protein to delineate the inner mitochondrial membrane (in red) using either a Nikon Eclipse 80i microscope (B) or an OMX 3D-SIM super-resolution imaging system (C and D). In D, intracellular parasites were treated for 8 hours with monensin (1 ng/mL). White arrowheads in D demarcate regions of Fis1 staining absent of the ATPase signal. Scale bar, 2 µm.

Since we had observed mitochondrial fragmentation following monensin treatment, we next sought to examine whether drug challenge resulted in altered localization of Fis1. RHΔ*hpt*+HAFis1 parasites were treated for 8 hours with monensin and prepared for super-resolution imaging. As anticipated, we observed fragmented mitochondrial morphology when examining the localization of F_1_B ATPase (Fig. 2D). Some of the mitochondrial fragments were encircled by the Fis1 protein, as one would expect for a protein in the OMM (Fig. 2D). Importantly, we also identified mitochondrial fragments that appeared to be connected by filaments of Fis1 (white arrowheads, Fig. 2D). These super-resolution images revealed that the OMM remains intact following the 8-hour monensin treatment despite the punctate appearance of the IMM. Thus, the observed effect of monensin treatment on mitochondrial morphology is not a true fragmentation but rather a constriction of the OMM in particular regions.

### The Fis1 transmembrane domain is required for proper localization to the OMM

Our previous studies have shown that the TM domain of Fis1 is sufficient for mitochondrial targeting (24). To determine whether the TM is necessary for mitochondrial localization we established a parasite strain expressing an exogenous copy of Fis1 with an N-terminal HA tag and truncated at the C-terminus as to lack the TM and CTS (Fig. 3A). Intracellular parasites of this strain were co-stained with antibodies against HA to detect Fis1ΔTM and against F_1_B ATPase to visualize the mitochondrion (Fig. 3B). Fis1 lacking the TM appears to be distributed throughout the cytoplasm in a punctate pattern (Fig. 3B). A similar result was observed when the TM of the endogenous Fis1 was replaced by an HA epitope tag using homologous recombination (Fig. 3C and D). Eliminating the TM of the endogenous Fis1 shifted its localization from the mitochondrion to the cytoplasm. Thus, proper Fis1 localization to the OMM is dependent on its C-terminal transmembrane domain and CTS.

**Figure 3.**
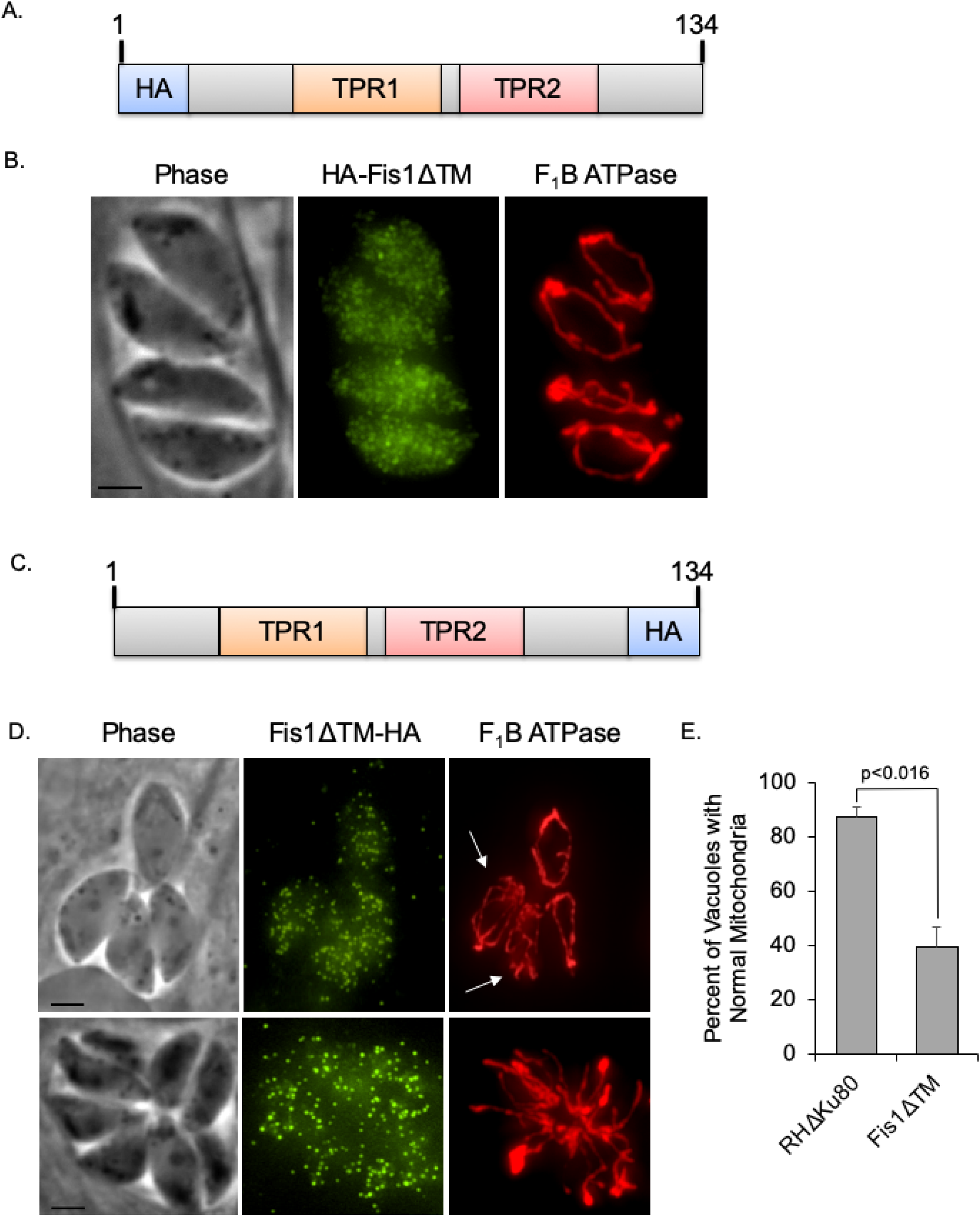
Fis1 localization is dependent on its transmembrane domain. To determine the necessity of the TM domain for localization of Fis1 we engineered strains in which either an exogenous or the endogenous Fis1 lacked the transmembrane domain. A. Schematic of the exogenous HA-FisΔTM. B. Parasites expressing HA-FisΔTM were co-stained for the exogenous Fis1 (in green) and the mitochondrial F_1_B ATPase (in red). Scale bar, 2 µm. C. Schematic of endogenous Fis1 in which TM has been replaced by an HA epitope (Fis1ΔTM-HA). D. Intracellular parasites of the strain expressing the truncated Fis1 were stained with antibodies against the HA tag (green) to detect Fis1ΔTM and antibodies against F_1_B ATPase (red) to detect mitochondria. White arrows indicate abnormal appearing mitochondria. Scale bar, 2 µm. E. The frequency of Fis1ΔTM-HA expressing parasites with abnormal mitochondrial morphology (extraneous fragments or branches) was examined and compared to that of the parental Δ*ku80* strain. In 3 independent experiments, parasite vacuoles from 15 random fields of view were enumerated, and the data are presented as percent of vacuoles with normal mitochondrial morphology ± SD. Student’s t-test was employed for determining statistical significance.

### Mitochondrial morphology is altered by the mislocalization of Fis1

When analyzing the localization of the truncated endogenous Fis1 we noted that the morphology of the mitochondrion appeared abnormal. Instead of the typical lasso seen in wildtype parasites (Figs. 1 and 2), the mitochondrion in parasites of the RHΔ*ku80*:Fis1ΔTM strain appeared to contain additional branches as well as unconnected strands, a phenotype that seemed to increase as the parasites underwent several rounds of division (Fig. 3D). In the RHΔ*ku80*:Fis1ΔTM strain, 60.4±7.5% of vacuoles had parasites with atypical mitochondrion (i.e. extraneous branches and strands). This is in contrast to the parental strain in which only 12.7±3.4% of vacuoles had parasites with atypical mitochondrion (Fig. 3D). These observations suggest that mislocalizing the endogenous Fis1 alters the typical mitochondrial morphology.

### RHΔ*ku80*:Fis1ΔTM parasites are less susceptible to monensin-induced mitochondrial disruption

The Fis1 protein in higher eukaryotes is responsible for fission of stressed and damaged mitochondria in order to maintain a healthy organelle pool. Thus, we next sought to determine the effect of Fis1 mislocalization in parasites undergoing monensin drug challenge. Parasites were vehicle treated or monensin treated for 12 hours. Cultures were fixed and examined by immunofluorescence microscopy and vacuoles with fragmented F_1_B ATPase signal were tallied. In the absence of any treatment the percentage of parasites with punctate F_1_B ATPase staining was statistically similar between parental and mutant strain (18.6±12.4% vs. 23.8±17.2). Following monensin treatment of parental parasites, an increase from 18.6±14.8% to 67.6±10.5% punctate mitochondria was observed for the parental strain (Fig. 4A). In RHΔ*ku80*:Fis1ΔTM parasites, an increase from 23.8±17.2 to 51.0±9.3% punctate mitochondria was recorded after drug challenge (Fig. 4A). The percent increase between the vehicle and monensin treated parasites was determined, and the increase in punctate mitochondria was statistically greater for the parental parasites compared to the RHΔ*ku80*:Fis1ΔTM parasites, 49.0±6.1% versus 27.2±9.6, respectively (Fig. 4B).

**Figure 4.**
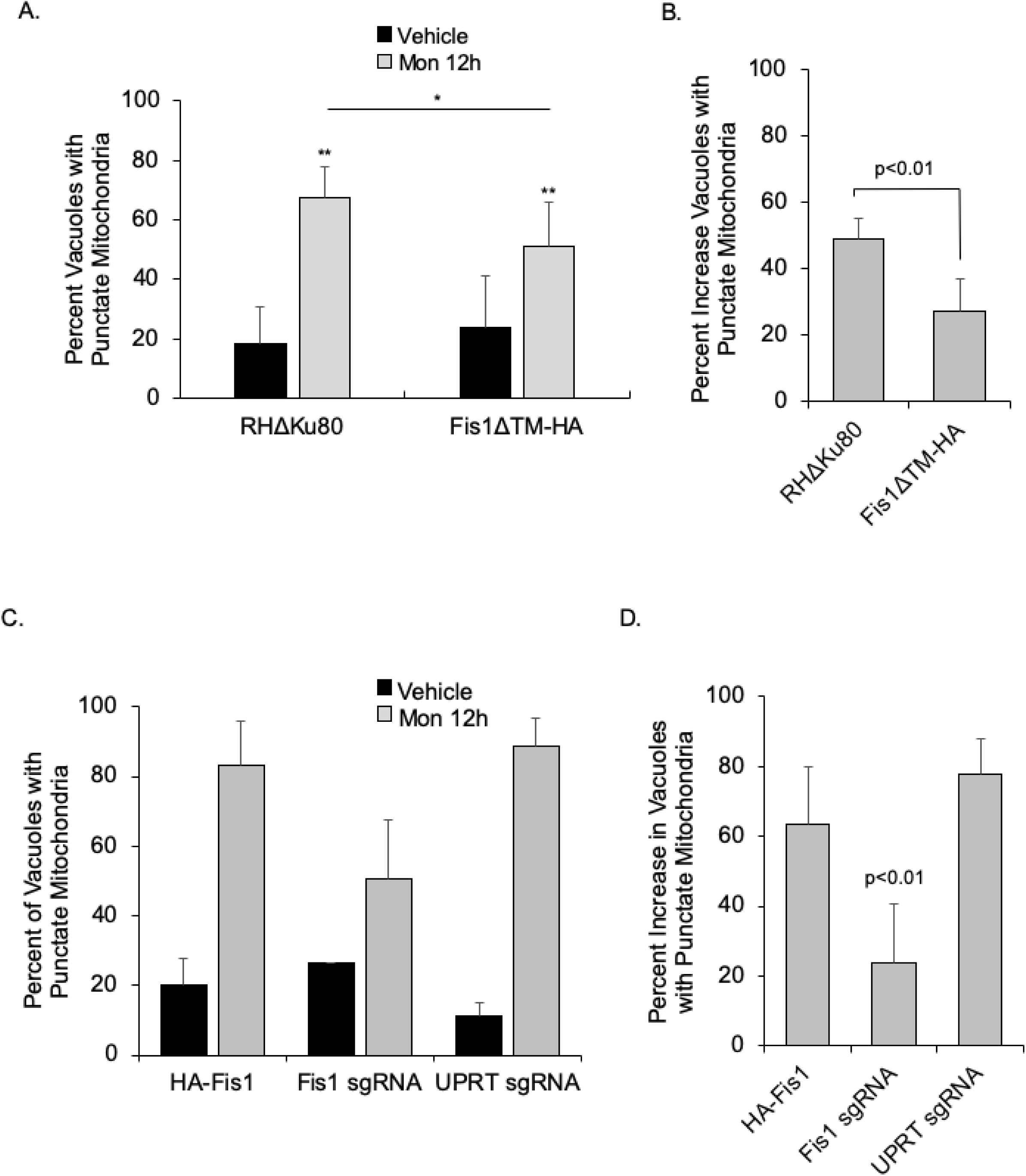
Disrupting Fis1 reduces monensin-induced mitochondrial remodeling. The ability of monensin to induce mitochondrial remodeling was assessed in strains expressing a mislocalized Fis1 or lacking Fis1. A. Parasites in which the endogenous Fis1 lacks the TM domain were vehicle or monensin treated for 12 hours. Parasite vacuoles were enumerated from 10 random fields of view for each strain and condition. The data are the average of 4 independent experiments and are presented as percent of vacuoles with punctate morphology ± SD. Statistical analysis was provided by one-way ANOVA post-hoc Tukey, where **p<0.001 as compared to vehicle. B. Data from A was analyzed to compare the number of vacuoles with punctate mitochondrion between untreated and treated parasites for each strain. Data is displayed as percent increase of vacuoles with punctate mitochondria upon treatment ± SD. C. RHΔ*hpt* parasites ectopically expressing the N-terminally HA tagged Fis1 were transfected with a plasmid expressing Cas9 and either a Fis1 specific sgRNA or the non-specific UPRT sgRNA. After transfection, parasites were immediately infected into HFFs on coverslips. Following approximately 16 hours in culture, cultures were vehicle or monensin treated for 12 hours and an IFA to monitor mitochondrial morphology was performed. The data presented are the averages of 6 coverslips from 2 independent transfections. Bars represent percent punctate mitochondria ± SD. Statistical significance was determined via Two-way ANOVA, treatment p<.0.0001, genotype p=0.006, genotype X treatment p=0.0003. D. The percent increase in punctate mitochondria between treatment and no treatment for the data shown in C was calculated and presented ± SD.

The phenotypes observed with the RHΔ*ku80*:Fis1ΔTM parasites could be due to either absence of Fis1 at the mitochondrion or a dominant negative effect from the mislocalized truncated protein. To differentiate between these possibilities, we next sought to determine how genetic ablation of Fis1 would affect the parasite’s ability to respond to monensin challenge and undergo mitochondrial remodeling. Employing the CRISPR/Cas9 system, RHΔ*hpt*+HAFis1 parasites ectopically expressing the N-terminally HA epitope-tagged Fis1 were either transfected with a sgRNA that would target both the endogenous and exogenous Fis1 gene or a sgRNA for the non-essential uracil phosphoribosyltransferase (UPRT) gene as a control. Parasites were immediately infected into human foreskin fibroblasts (HFFs) on coverslips and grown in culture for 16 hours. Infected cultures were then vehicle or monensin treated for 12 hours. After treatment parasites were fixed and IFA was performed staining for HA to detect Fis1 and F_1_B ATPase to visualize the mitochondrion (Supplemental figure S1). We then compared the mitochondrial morphology in Fis1 sgRNA transfected parasites lacking HA signal to that of control sgRNA transfected parasites with HA signal. To control for the effects of Cas9, which is fused to GFP (25), we only analyzed the mitochondrial morphology in parasites with nuclear GFP signal. In vehicle treated parasites, there was no significant difference between parasites lacking HA-tagged Fis1 expression and those targeted for the UPRT when compared to the control parasites still expressing Fis1 (Fig. 4C and Supplemental figure S1). Interestingly, in contrast to what we observed with mislocalized Fis1, complete lack of Fis1 did not affect the mitochondrial morphology. Thus, it appears that Fis1 lacking the TM domain imparts a dominant negative effect on mitochondrial morphology.

As expected, monensin treatment of the Cas9/sgRNA transfected parasites resulted in an increase in the number of vacuoles containing punctate F_1_B ATPase signal. Control non-transfected parasites and those transfected with the UPRT sgRNA possessed 83.3±12.8% and 88.8±7.7% punctate mitochondria, respectively (Fig. 4C). However, parasites lacking the HA-tagged Fis1 signal displayed significantly fewer vacuoles with disrupted mitochondria upon monensin treatment, comprising only 50.5±16.9% of the vacuoles (Fig. 4C). The percent increase in the number of vacuoles with punctate mitochondria for the parasites deficient in HA-tagged Fis1 expression was significantly lower than for either control parasite populations, 23.9±17% versus 63.3±16.8% and 77.8±10.2%, respectively (Fig. 4D). Two-way Anova analysis indicated that there was both a treatment and genotype effect and that the effects interact. Overall, these data indicate that complete lack of Fis1 does not affect mitochondrial morphology in untreated parasites but significantly decreases monensin-induced mitochondrial remodeling, indicating that Fis1 is partially required for constriction of the IMM in response to treatment with the ionophore.

### A putative Fis1 interactor localizes to the OMM

Mislocalization of the endogenous Fis1 results in a dominant negative phenotype in terms of mitochondrial morphology. We hypothesize that this is the result of mislocalization of Fis1 interactors required at the mitochondrion for normal morphology. To identify these potential interactors, we employed a Yeast Two-Hybrid (Y2H) interaction screen. Using full-length Fis1 as bait, 46 million clones were screened for Y2H interaction and 247 were selected for identification. The putative interactors were then given a confidence score based on the likelihood of interaction with Fis1 (26, 27). This resulted in 24 potential interactors with a global Predicted Biological Score (PBS) from A (highest confidence) to D (lowest confidence) (26, 27) (Table 1). To narrow down the list we immunoprecipitated the exogenous HA tagged Fis1 using HA conjugated beads and analyzed the precipitated complex by mass spectroscopy. As a control, we used the parental RHΔ*hpt* strain, which does not express the hemagglutinin tag. Through this analysis, we identified 11 putative interactors that had at least 5 peptides in the Fis1 sample and no peptides in the control sample (Table S1 in supplemental material). Among these only one was also identified in the Y2H interaction screen, TGGT1_265180.

**Table 1.** Proteins identified as Fis1 interactors through a yeast two hybrid screen. Predicted biological scores (PBS) are confidence score, with A indicating the highest confidence of interaction and D being the lowest (14). Highlighted are proteins also identified in the mitochondrial proteome (41).

| Gene ID | Product Description | PBS |
| --- | --- | --- |
| TGGT1_215520 | hypothetical protein | A |
| TGGT1_218560 | acetyl-coA carboxylase ACC2 | B |
| TGGT1_222800 | glycogen synthase, putative | B |
| TGGT1_265180 | hypothetical protein | B |
| TGGT1_224270 | hypothetical protein | C |
| TGGT1_293840 | hypothetical protein | C |
| TGGT1_201390 | hypothetical protein | D |
| TGGT1_226050 | hypothetical protein | D |
| TGGT1_237015 | GRA43 | D |
| TGGT1_246720 | hypothetical protein | D |
| TGGT1_247700 | AP2 domain transcription factor AP2XII-4 | D |
| TGGT1_284620 | hypothetical protein | D |
| TGGT1_286470 | AGC kinase | D |
| TGGT1_287980 | FHA domain-containing protein | D |
| TGGT1_297770 | hypothetical protein | D |
| TGGT1_299670 | hypothetical protein | D |
| TGGT1_304990 | guanylate-binding protein | D |
| TGGT1_321370 | hypothetical protein | D |
| TGGT1_321450 | Myb family DNA-binding domain-containing | D |

To determine the localization of TgGT1_265180, we introduced a C-terminal myc epitope tag to the endogenous gene. IFA assays of the resulting strain show that, like Fis1, TgGT1_265180 is localized to the mitochondrion of intracellular parasites (Fig. 5A). This association with the mitochondrion persists during parasite division (Fig. 5B). To determine whether the protein is associated with the outside or inside of the mitochondrion we performed IFA after permeabilization with various concentrations of digitonin using detection of F_1_B ATPase to monitor mitochondrial permeabilization (Fig. 5C). When using 0.01% digitonin we can detect both F_1_B ATPase and TgGT1_265180 (Fig. 5C). By contrast, using 0.005% digitonin allows for detection of TgGT1_265180 but not F_1_B ATPase, which suggest that TgGT1_265180 likely associates with the OMM and faces the cytoplasm of the parasite (Fig. 5C). Association with the OMM was confirmed by treatment with monensin. After treating TgGT1_265180(myc) expressing parasites with monensin, we observed a similar pattern to that of Fis1 in which fragments containing the IMM marker F_1_B ATPase are surrounded and connected by TgGT1_265180 (Fig. 5D). Thus, TgGT1_265180 localizes to the OMM as expected for a *bona fide* interactor of Fis1.

**Figure 5.**
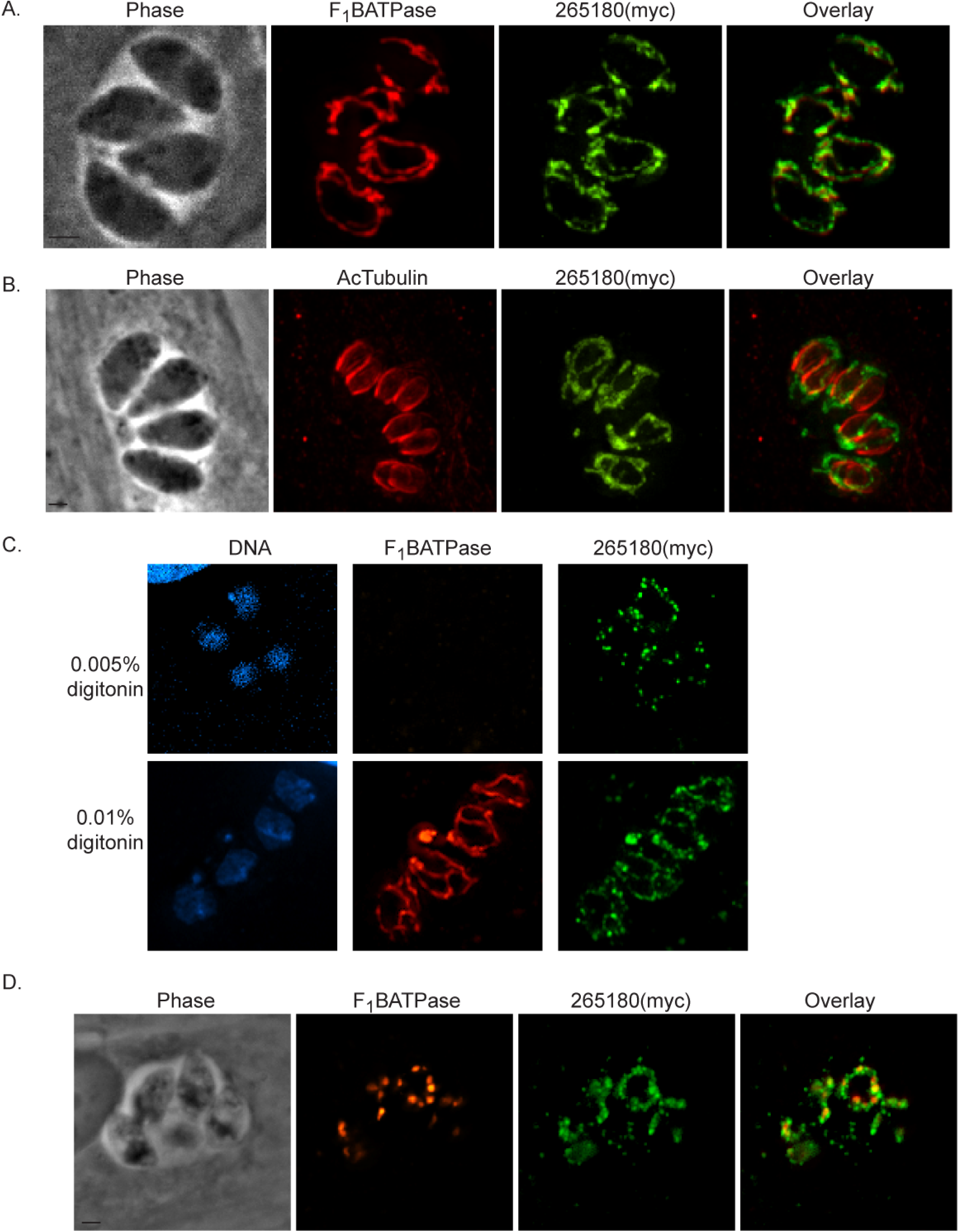
Fis1 interactor TgGT1_265180 localizes to the outer mitochondrial membrane. To investigate the localization of TgGT1_265180 we introduced sequences encoding an N-terminal myc tag to the endogenous locus. A. Intracellular parasites of the TgGT1_265180(myc) expressing strain were stained for the mitochondrial F_1_B ATPase (red) and for myc (green). B. Intracellular parasites of the same strain were stained for myc (green) and acetylated tubulin (red), which clearly demarcates daughter parasites during division. C. Intracellular parasites of the TgGT1_265180(myc) expressing strain were fixed and permeabilized with either 0.005% or 0.01% digitonin before staining for the IMM protein F_1_B ATPase (red) and myc (green). TgGT1_265180 can be detected when F_1_B ATPase remains inaccessible to the antibodies suggesting that it is associated with the OMM. D. TgGT1_265180(myc) parasites were treated with 5 mM monensin for 5 hours. Mitochondrial morphology was monitored by IFA for TgGT1_265180(myc) (green) and F_1_B ATPase (red). Scale bar, 2 µm.

### Localization of TgGT1_265180 is partially dependent on proper Fis1 localization

Despite its association with the OMM, TgGT1_265180 has no predicted trans-membrane domains or posttranslational modifications that would suggest membrane interaction. Therefore, we hypothesize the localization of TgGT1_265180 occurs via protein-protein interaction. To test this idea, we transfected parasites with an ectopic copy of either full length or truncated TgGT1_265180 carrying a C-terminal HA epitope tag and under the control of the TgGT1_265180 promoter (Fig. 6A). The truncated form lacks the C-terminal 92 amino acids, which represent the region of the protein that was identified through the Y2H screen as interacting with Fis1, referred to as the Selected Interaction Domain (SID). As expected, the full-length ectopic copy localized to the mitochondrion (Fig. 6A). However, deletion of the SID resulted in the mislocalization of the protein to the cytoplasm (Fig. 6A). These data indicate that the C-terminal SID is necessary for proper mitochondrial localization.

**Figure 6.**
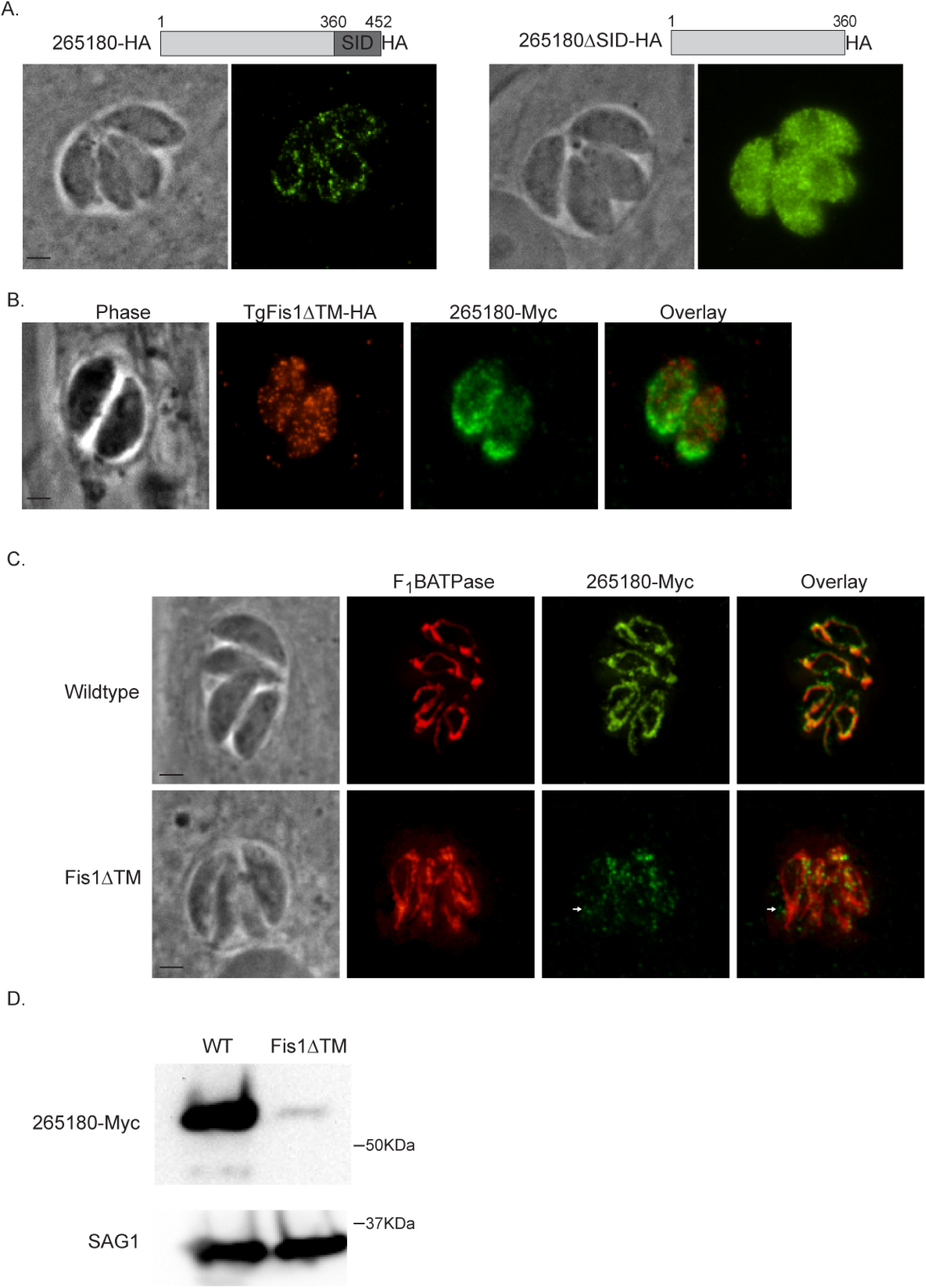
Association of TgGT1_265180 with the mitochondrion depends on Fis1. To investigate how TgGT1_265180 associates with the mitochondrion we tested the role of its C-terminus and of Fis1 on its localization. A. Parasites were transfected with an exogenous copy of C-terminally HA tagged wildtype TgGT1_265180 or with N-terminally HA tagged TgGT1_265180 lacking the Selected Interaction domain (SID). The SID is the region of TgGT1_265180 that was identified as interacting with Fis1. Intracellular parasites expressing TgGT1_265180-HA (left) or TgGT1_265180ΔSID-HA (right) were stained for HA. B. Intracellular FisΔTM-HA parasites expressing an endogenous copy of C-terminally myc tagged TgGT1_265180 were probed for HA to detect Fis1 (red) and for myc to detect TgGT1_265180 (green) C. Wildtype or FisΔTM-HA parasites endogenously expressing TgGT1_265180-Myc were stained for F_1_B ATPase (red) and myc (green) to monitor localization of TgGT1_265180. Scale bar, 2 µm. D. Representative Western blot of extract from wildtype (WT) and Fis1ΔTM parasites expressing TgGT1_265180-myc probed for myc (top blot) and for SAG1 (bottom blot) as a loading control.

To investigate if localization of TgGT1_265180 to the mitochondrion is through an interaction with Fis1, we added a myc epitope tag to the endogenous TgGT1_265180 in the strain in which Fis1 lacks its TM (RHΔ*ku80*:Fis1ΔTM) and is mislocalized to the cytoplasm. In this strain, TgGT1_265180 does not colocalize with the mislocalized Fis1 but appears to accumulate towards the basal end of the parasites in a pattern that does not resemble normal mitochondrial localization (Fig. 6B). To further analyze the localization of TgGT1_265180 in the RHΔ*ku80*:Fis1ΔTM parasite line, we co-stained for F_1_B ATPase (Fig. 6C). While we observed some overlap between the TgGT1_265180 and F_1_B ATPase signals, TgGT1_265180 was also detected away from the mitochondrion (Fig. 6C). Interestingly, we observed that the TgGT1_265180(myc) signal, as detected through IFA, appeared to be much weaker in the Fis1ΔTM strain than in the parental one (Fig. 6C). To quantitate this observation, we performed Western blots from both strains probing for TgGT1_265180(myc) (Fig. 6D). This analysis corroborated that indeed the levels of endogenous TgGT1_265180 are significantly reduced when Fis1 is mislocalized away from the mitochondrion (Fig. 6D). We quantitated the levels of TgGT1_265180 in both strains with densitometry of three independent Western blots using the surface antigen SAG1 as a loading control and determined that the level of TgGT1_265180 in the RHΔ*ku80*:Fis1ΔTM is 23.2±8.7% of that in the parental strain. In conjunction, these results indicate that TgGT1_265180 associates with the mitochondrion via its C-terminus and that its localization and stability is at least in part dependent on Fis1.

### 265180 knockout affects parasite fitness in tissue culture

Based on a genome-wide CRISPR screen, TgGT1_265180 was assigned a relative fitness phenotype score of −1.65, which indicates that, while its absence would negatively affect parasite fitness, it is likely not essential, making its genetic disruption possible (28). Accordingly, we employed double homologous recombination to replace the coding sequence of TgGT1_265180 with a drug selection marker (Fig. 7A). Proper integration of the knockout construct in stably transfected clones was confirmed using PCR (Fig. 7B). To test the effect of the knockout on parasite propagation we used a standard growth assay in which the same number of either parental or mutant parasites were allowed to infect human fibroblasts and form plaques over a five-day period. We observed a significant propagation defect in the *Δ265180* parasites, exhibited by both less and smaller plaques in comparison to the parental strain. To quantitate this defect, we counted the number of plaques formed by the parental and knockout strains in three separate experiments each with experimental triplicates (Fig. 7C). The average number of plaques by the *Δ265180* was 30.2±9.0% of that detected for the parental strain.

**Figure 7.**
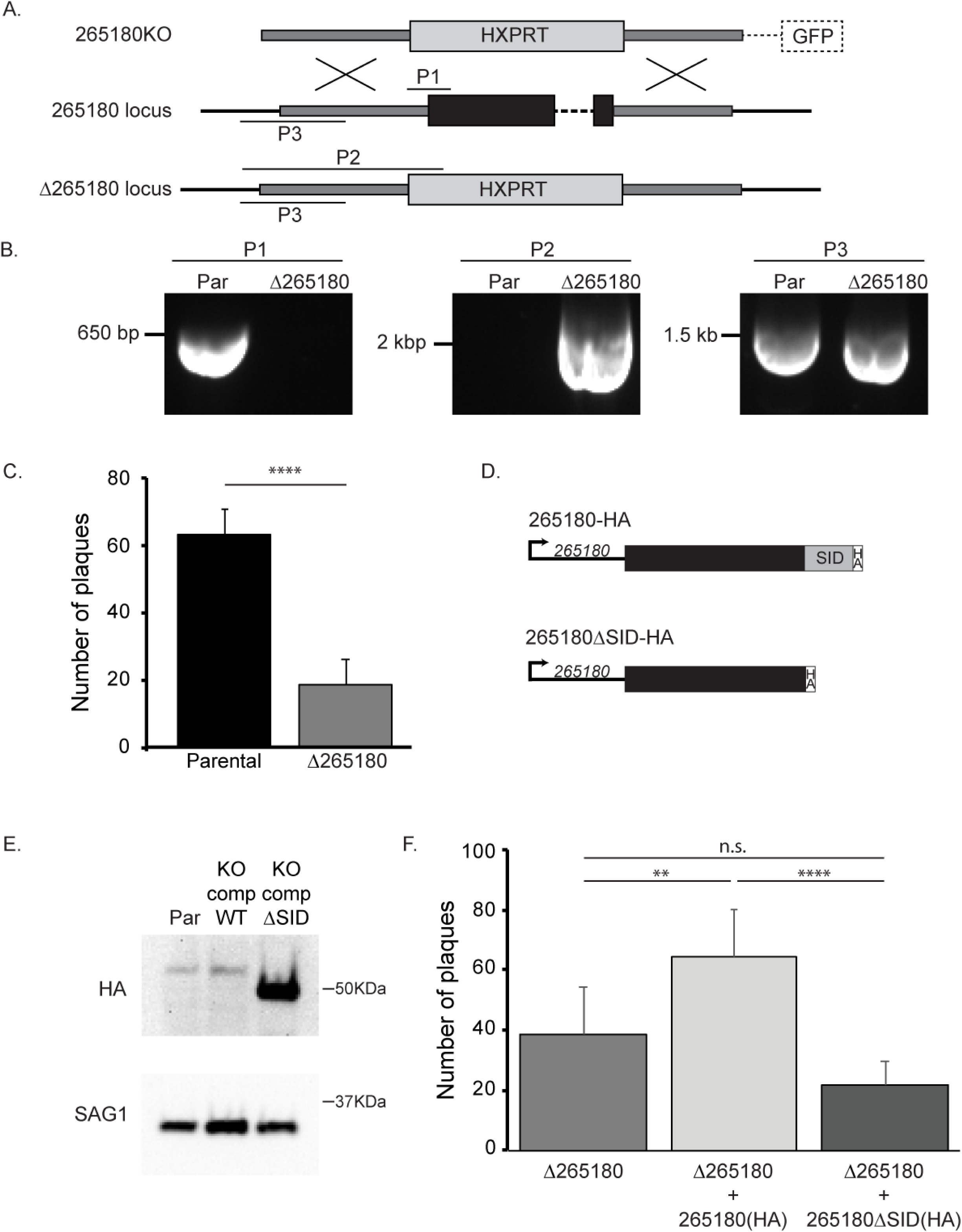
Knockout of TgGT1_265180 affects parasite propagation. To investigate the role of TgGT1_265180 in parasite fitness we established knockout and complemented strains. A. Schematic of strategy implemented to disrupt the TgGT1_265180 by replacing the coding sequences by the selectable marker *HPT*. On top is the vector used to drive the gene replacement, which includes *HPT* flanked by areas of homology to the TgGT1_265180 locus (dark grey boxes) and a downstream copy of GFP that is not integrated upon the desired double homologous recombination and can be used as a negative selectable marker. Endogenous TgGT1_265180 is depicted in the middle with coding sequences represented by a black box. Bottom drawing shows the expected result from gene replacement in the knockout strain. P1, P2, and P3 indicate the PCR amplicons that were used to confirm integration. P1 would only be detected from parental parasites, P2 only from knockout parasites and P3 from both. B. PCR products from reactions to detect P1, P2 and P3 in the parental strain and the established Δ*265180* clone. C. Average number of plaques per well for either parental or knockout strains after 4-day incubation period. Plaque assays were done in biological and technical triplicates, with error bars representing ±SD. Statistical analysis via t-test, ****p<0.0001 D. Diagrams depict the two constructs used for complementation: TgGT1_265180-HA and TgGT1_265180ΔSID-HA. SID is the Selected Interaction Domain identified through the two-hybrid screen. E. Representative Western blot of a strain in which the endogenous TgGT1_265180 includes a HA epitope tag (Par), and the knockout strain complemented with wildtype TgGT1_265180-HA (KO comp WT) or with TgGT1_265180ΔSID-HA (KO comp ΔSID) probed for HA (top blot) and for SAG1 (bottom blot) as a loading control. F. Average number of plaques per well for each strain after 4-day incubation period. Plaque assays were done in biological and technical triplicates, with error bars representing ±SD. Statistical analysis performed using One-way Anova, ****p<0.0001 and **p<.0019

To confirm that the phenotype observed was due to the disruption of the target gene and not a secondary effect, we complemented the *Δ265180* strain with an exogenous copy of the TgGT1_265180 cDNA including a C-terminal HA epitope tag and driven by its own promoter. As the knockout strain lacks Ku80 and does not effectively allow for random integration, the exogenous copy was directed to the remnants of the Ku80 locus using CRISPR/Cas9. In addition to complementing with the wildtype TgGT1_265180, we transfected the knockout strain with the truncated version TgGT1_265180ΔSID, which does not localize to the mitochondrion (Fig. 7D). Western blot showed that both complemented strains expressed proteins of the expected size (Fig. 7E). Interestingly, while the wildtype complement expression level is similar to that of the endogenous levels, the truncated copy appears to be expressed at a much higher level (Fig. 7E). Plaque assays of both the *Δ265180*+265180(HA) and *Δ265180*+265180ΔSID(HA) strains were performed in parallel to the knockout strain (Fig. 7F). The average number of plaques by the *Δ265180*+265180(HA) was 64.5±15.8, which is significantly higher than both the knockout and truncated complement strains (Fig. 7F). *Δ265180*+265180ΔSID(HA) had a lower average number of plaques (21.6±8.0) than that of the knockout (38.8±15.3), but this difference was not statistically significant. These results indicate that proper localization of TgGT1_265180 is necessary to rescue the growth phenotype seen in tissue culture.

### 265180 disrupts the normal morphology of the mitochondrion

As TgGT1_265180 is associated with the mitochondrion we assessed mitochondrial morphology in the knockout parasites. In intracellular parasites, the mitochondrion maintains what is referred to as a lasso shape that abuts the periphery of the parasite (17) (Figs. 2 and 8A). However, based on staining with antibodies against F_1_B ATPase, the mitochondrion of *Δ265180* parasites exhibit an altered mitochondrial morphology, with the bulk of the mitochondrial material concentrated at one end of the parasite (Fig. 8A). By contrast disruption of TgGT1_265180 did not affect the morphology of the apicoplast, rhoptries, or endoplasmic reticulum (Fig S2). Introduction of the wild type TgGT1_265180 to the knockout strain complements the mitochondrial phenotype (Fig. 8B). In contrast, the truncated TgGT1_265180*Δ*SID, which is not localized to the mitochondrion, does not rescue the collapsed mitochondrion phenotype (Fig. 8C). The phenotype of the knockout and the complemented strains was quantitated by determining the percentage of parasites with normal and abnormal mitochondrion morphology. Normally, the *Toxoplasma* mitochondrion retracts from the periphery of the parasite during egress and changes its morphology to what has been described as sperm-like and collapsed (17). Interestingly, we observed all three morphologies normally associated with extracellular parasites (lasso, sperm-like, and collapsed) in intracellular parasites of the *Δ265180* strain (Fig. 9A). With the parental strain, the proportion of mitochondrial morphologies in intracellular parasites is 84.7±2.1% lasso, 15.3±2.1% sperm-like, and 0% collapsed. By contrast, intracellular parasites of the *Δ265180* strain, the mitochondrial distribution is 6.0±2.6% lasso, 50.0±2% sperm-like, and 44.0±4.4% collapsed (Fig. 9B). Just as it was the case for the plaquing phenotype, introduction of a wild type copy of TgGT1_265180 partly rescues the morphological phenotype with 48.5±4.4% of parasites exhibiting lasso-shaped mitochondrion, 49.2±3.9% sperm-like, and only 2.3±0.6% collapsed. Additionally, the truncated copy had a similar morphological distribution to that of the knockout strain (2.7±2.3% lasso, 56.0±10.1% sperm-like, and 41.4±12.4% collapsed) and was significantly different from the distributions of the parental and complement strains, which is consistent with defects seen in plaquing (Fig. 9B). Thus, TgGT1_265180 plays a crucial role in maintaining proper morphology of the mitochondrion. Consequently, we have dubbed this new gene Lasso Maintenance Factor 1 (LMF1).

**Figure 8.**
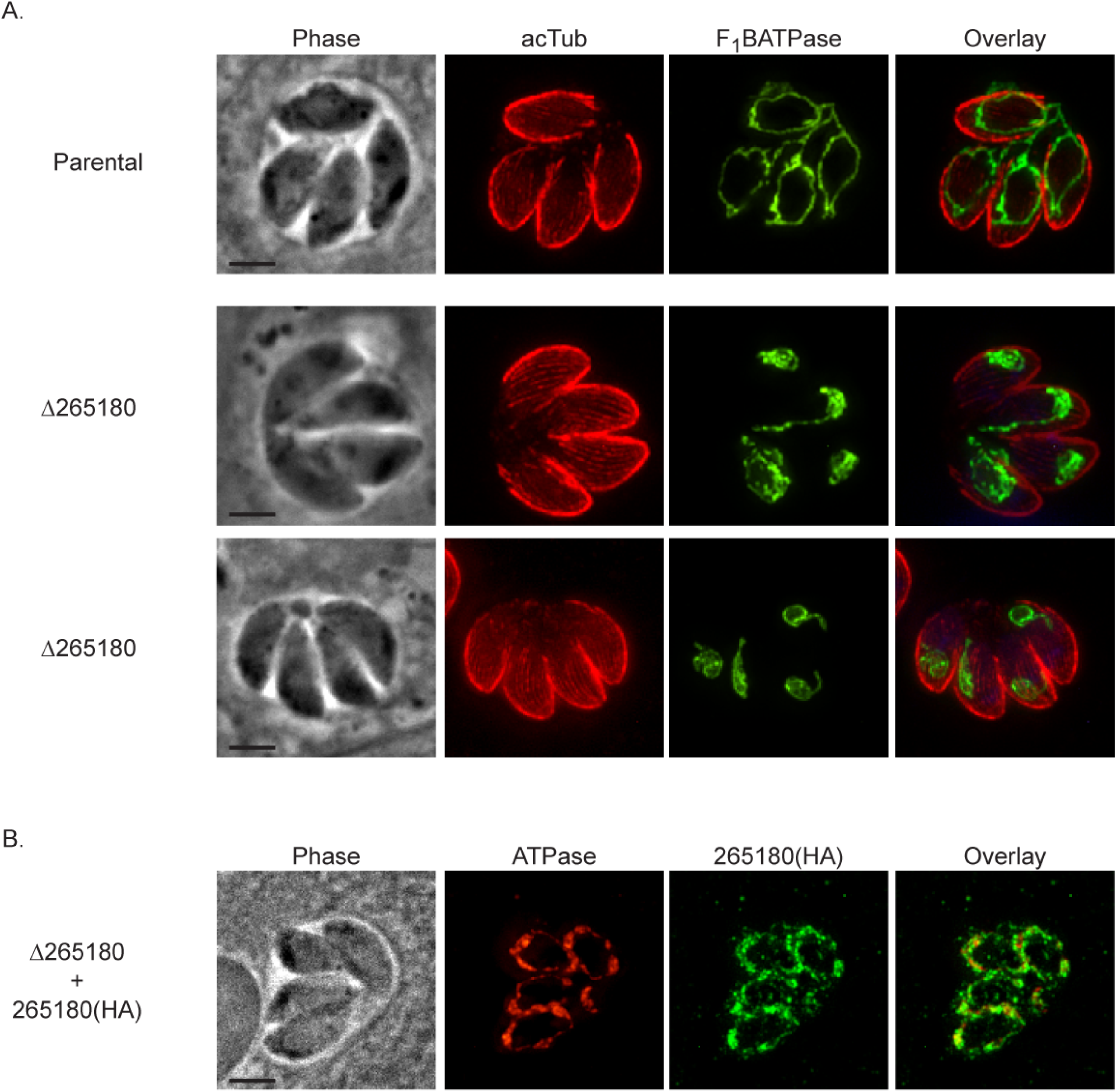
Mitochondrial morphology is disrupted by lack of TgGT1_265180. To determine the effect of TgGT1_265180 ablation on the mitochondrion knockout and complemented parasites were analyzed by IFA. A. Intracellular parasites of the parental or the Δ*265180* strain were stained for F_1_B ATPase (green) to monitor mitochondrion and for acetylated tubulin (acTub) to detect the parasite cytoskeleton (red). B. and C. IFA of knockout parasites (Δ2*65180*) transformed with either the wildtype (265180(HA)) or truncated TgGT1_265180 (265180ΔSID(HA)) with antibodies against F_1_B ATPase (red) and HA (green). Scale bar, 2 µm.

**Figure 9.**
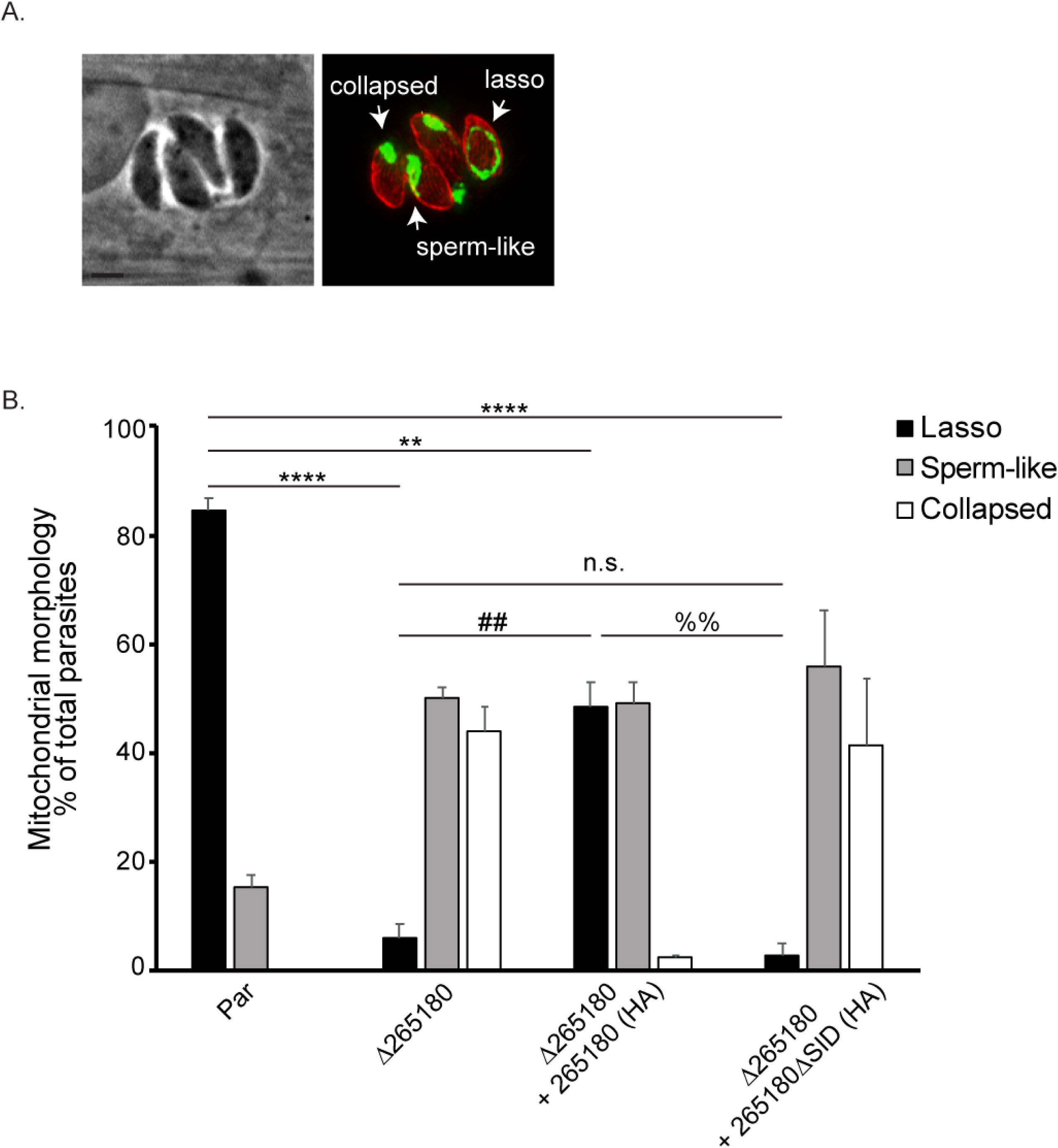
Intracellular parasites lacking TgGT1_265180 do not maintain their mitochondrion in the lasso conformation. To determine the penetrance of the mitochondrial phenotype observed in with the Δ*265180* strain the different morphological patterns observed were quantitated. A. Intracellular parasites of the Δ*265180* stained for F_1_B ATPase (green) and acetylated tubulin (red) exhibiting three distinct mitochondrial morphologies: lasso, collapsed, and sperm-like. Scale bar, 2 µm B. Percentage of parasites with each of the three different morphologies for the parental (par), knockout (Δ*265180*) and complemented strains (Δ*265180*+265180(HA) and Δ*265180*+265180ΔSID(HA)). Data is average of biological triplicates, at least 50 vacuoles per sample were inspected. Error bars are SD. Statistics shown are ANOVA of percentage of parasites with lasso shape for each strain. ****p<0.001, **p<0.004, ##p<0.003, and %%p<0.002.

### Disruption of LMF1 results in defects in mitochondrial segregation between daughter parasites

During our analysis of mitochondrial morphology in the LMF1 mutant strain we noted various aberrant phenotypes that likely relate to parasite and mitochondrial division. *Toxoplasma* divides through a process called endodyogeny, where two daughter parasites form within a mother parasite (29). This results in a doubling in the number of parasites in a vacuole after each round of replication. We noted that vacuoles of the LMF1 strain often had abnormal number of parasites (i.e. not 2, 4, 8, etc). We found that approximately 25.3±5.1% of vacuoles in *Δ265180* parasites had odd numbers compared to 5.8±2.9% in wildtype parasites and 13.7±3.1% in the complemented strain (Fig. 10A). Interestingly, we also noticed numerous vacuoles in which some parasites lacked a mitochondrion based on absence of F_1_B ATPase staining (Fig. 10B, white arrows). When quantified, 16.2±4.0% of vacuoles contained at least one parasite that did not have mitochondrial material compared to 0.3±0.6% of RHΔ*ku80* parasites were amitochondriate (Fig. 10B). As with the other phenotypes, exogenous expression of wildtype LMF1 complemented the phenotype with 6.0±1.7% of vacuoles containing amitochondriate parasites. In addition to amitochondriate parasites, disruption of LMF1 also results in an accumulation of mitochondrial material outside of parasites (Fig. 10C, white arrows). We determined that 30.9±4.0% of vacuoles had extraparasitic mitochondrial material, which is three times greater than that of the parental parasite line (10.6±3.2%). Interestingly, this particular phenotype was not complimented, as 28.3±2.1% of *Δ265180*+265180(HA) vacuoles contained extraparasitic material (Fig. 10C).

**Figure 10.**
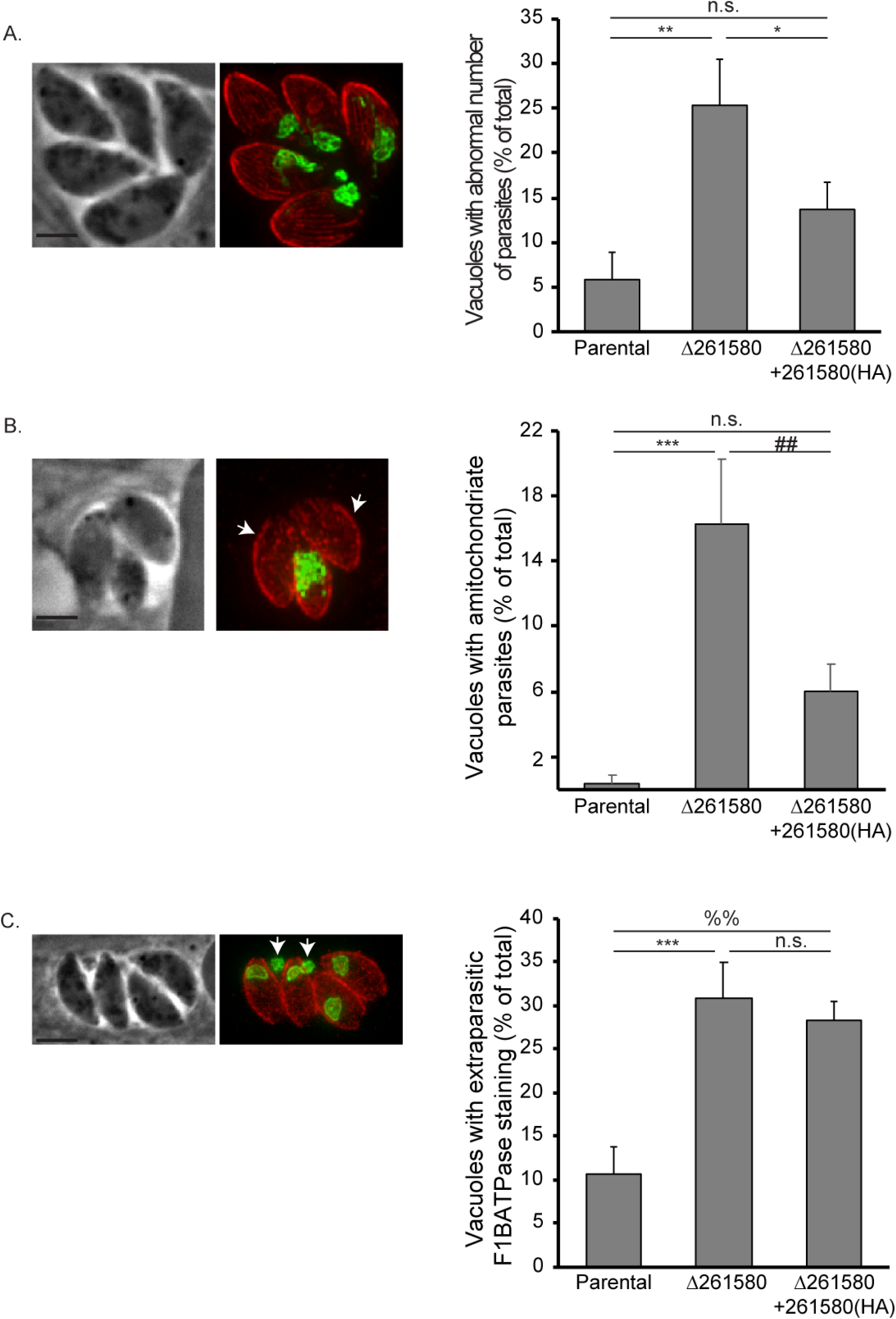
Parasites lacking TgGT1_265180 exhibit various division related phenotypes. IFA of knockout parasites stained for F_1_B ATPase (green) and acetylated tubulin (red) reveal various aberrant phenotypes. A. Image on the left is of Δ*265180* vacuole containing five parasites rather than either four or eight as expected. Graph shows the percentage of vacuoles with abnormal number of parasites for the three strains. B. Image shows vacuole with amitochondriate parasites (arrows) based on lack of F_1_B ATPase signal. Graph shows the percentage of vacuoles with at least one amitochondriate parasite for each strain. C. Image is of vacuole that contains parasites with F_1_B ATPase signal outside of the parasite and within the parasitophorous vacuole (arrows). Scale bar, 2 µm. Graph shows the percentage of vacuoles with this phenotype. For all graphs n=3 ±SD with at least 50 vacuoles per sample inspected. Statistical analysis done with on-way ANOVA Tukey post-hoc, ***p<0.0006 **p<002, *p<.02, ##p<0.006, %%p<0.001

We hypothesize that these phenotypes (abnormal number of parasites, amitochondriate parasites, and extraparasitic mitochondria) are the result of aberrant segregation of the mitochondrion into the daughter cells during endodyogeny. Accordingly, we co-stained parental and knockout parasites for acetylated tubulin to detect daughter cells and for F_1_B ATPase to monitor the mitochondrion (Fig. 11). During the early (E) stages of division, wildtype parasite mitochondria surround the forming daughters (Fig. 11A top panel). As endodyogeny progresses to an intermediate (I) stage, the mitochondrion remains excluded from the daughters (Fig. 11A middle panel). When the daughters have almost fully formed (late (L) stages), branches of mother mitochondria incorporate into the daughter parasites before emerging from the mother (Fig. 11A bottom panel). When LMF1 is disrupted, the mitochondrion does not have the typical lasso shape and appears to associate with one of the two daughters instead of surrounding both (Fig. 11B top panel, E). As the daughters continue to form in the LMF1 deficient parasites, the mitochondrial material remains associated with one daughter or is completely excluded from the budding daughters (Fig. 11B second panel, I). During the final stages of endodyogeny, some daughters seem to have received mitochondrial material, whereas others have not. This correlates to an accumulation of mitochondrial material outside of the parasites (Fig. 11B bottom three panels, L). Therefore, disruption of LMF1 leads to defects in mitochondrial segregation during endodyogeny, which agrees with the aberrant phenotypes observed with mitochondrial shape and localization (Figs. 9 and 10).

**Figure 11.**
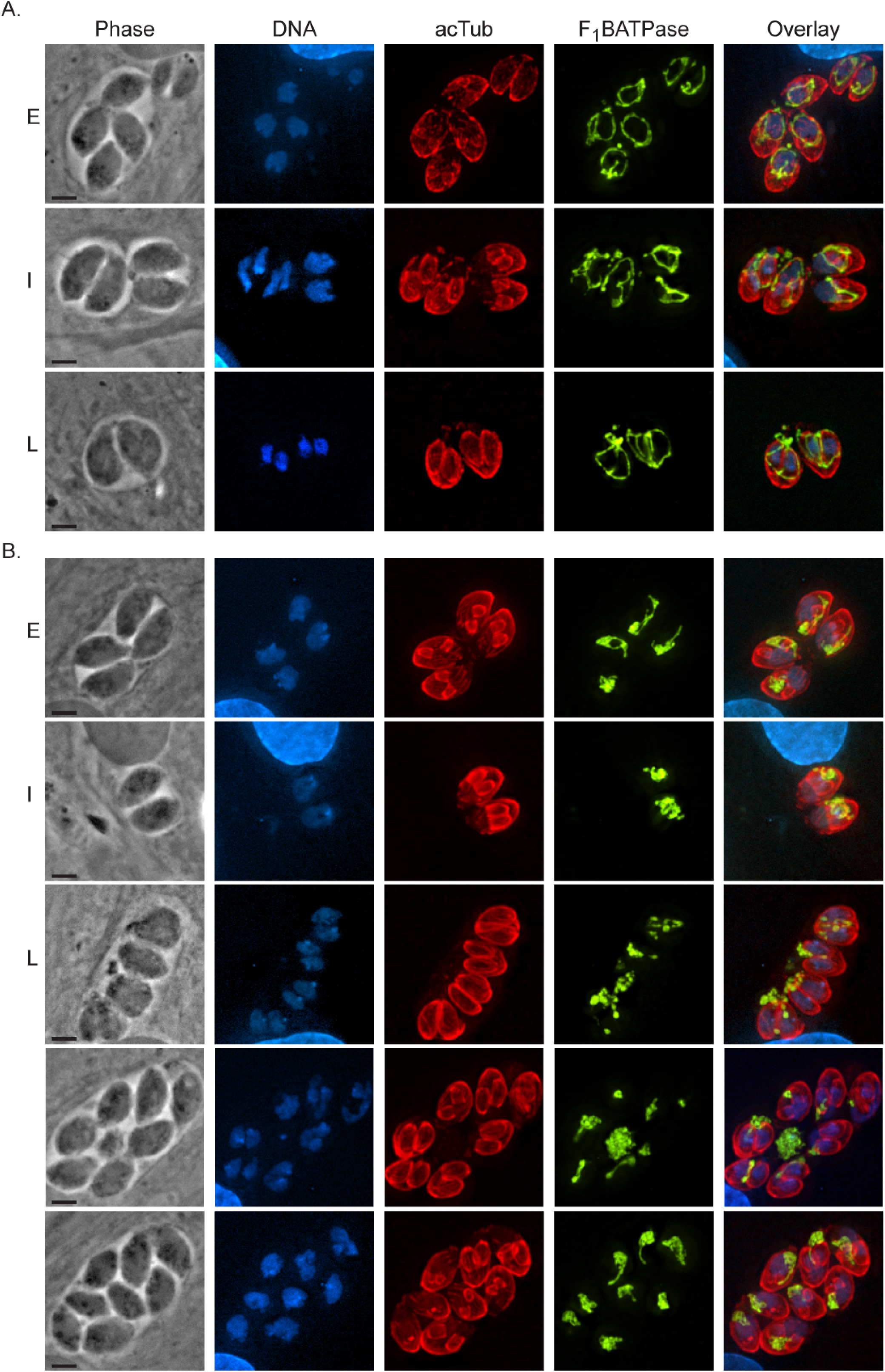
TgGT1_265180 disruption results in mitochondrial segregation defects. To examine mitochondrial dynamics during parasite division IFAs of parasites during early (E), intermediate (I), and late (L) stages of endodyogeny were conducted. A. IFAs of intracellular wildtype parasites. B. IFAs of intracellular Δ*265180* (aka LMF1) knockout parasites. In both A and B stage of division was determined by DAPI staining (blue) and acetylated tubulin (red), which demarcate budding daughters. Mitochondrial morphology was observed by staining with F_1_B ATPase, shown here in green. Scale bar, 2 µm.

## DISCUSSION

The single mitochondrion of the pathogen *Toxoplasma gondii* is highly dynamic, with its location and structure changing during various stages of the parasite’s lytic cycle. As the last organelle to move from a live mother parasite into two nascent daughter cells, the morphology and position of the mitochondrion is tightly regulated during parasite division. Similarly, as the parasite moves from inside to outside host cells the mitochondrion morphology dramatically changes. While inside the host cell *Toxoplasma*’s mitochondrion forms a lasso with multiple points of contact with the parasite pellicle, then quickly retracts from the parasite periphery to a collapsed bundle at the apical end as the parasites move to the extracellular space. In this study, we show that the mitochondrial morphology also changes under treatment with the anti-parasitic drugs atovaquone and monensin. Under drug treatment the mitochondrion’s outer membrane becomes constricted causing the inner mitochondrial material to appear punctate. Importantly, this phenomenon is completely reversible and upon removal of monensin the mitochondrion returns to its typical shape. We also show that mitochondrial constriction upon monensin treatment is in partly dependent on the presence of the fission protein Fis1 at the mitochondrion. Thus, we have discovered a mechanism by which the parasite reversibly restructures its mitochondrion.

The morphological changes experienced by the mitochondrion under monensin treatment are likely a response to stress and might represent a mechanism by which the parasite protects the mitochondrion from irreversible damage. Mitochondria from numerous organisms alter their morphology to respond to specific stressors, such as UV radiation and nutrient starvation (30–33). In conditions that damage mitochondrial DNA, such as cycloheximide and UV radiation, mitochondria hyperfuse (30). This phenomenon most likely occurs to complement damaged mitochondrial DNA and promote DNA mixing, and is dependent on mitochondrial fusion factors such as OPA1 and Mfn1/2 (30). Conditions that affect mitochondrial respiration, such as oligomycin and uncoupling agents, cause mitochondrial fragmentation (32, 33). Nutrient conditions also play a role in mitochondrial morphology. For example, yeast cultured in aerobic, respiratory conditions have more punctate mitochondria whereas anaerobic conditions result in branched and elongated morphologies (31). The smaller, more punctate mitochondria have higher surface area than those with branched morphology, indicating a higher respiratory capacity (31). Constriction of the inner mitochondrial membrane is a priming event for mitochondrial division and can be augmented by changes in Ca^2+^ levels, producing a “beads-on-a-string” phenotype similar to that observed with monensin. These data suggest that mitochondrial morphology is dependent upon environmental conditions and stressors. Therefore, the phenotype we see under monensin treatment is likely a protective mechanism for the mitochondrion against the effects of the ionophore.

As the effect of monensin is a reversible constriction along the outer mitochondrial membrane, we hypothesize that this phenomenon would require the mitochondrial fission machinery. The yeast mitochondrial fission machinery is the most well characterized and it is comprised of the membrane anchored protein Fis1p, which actively recruits other proteins to the mitochondria during fission like Mdv1 (mitochondrial division protein 1), which acts as an adapter protein. Fis1p is then able to recruit a GTPase, dynamin (Dmn1), which is able to drive the final scission of the mitochondrion (11). No homologs for Mdv1 have been found in *Toxoplasma gondii*, but there are one Fis1 homolog (TGGT1_263323) and three dynamin-related proteins: DrpA, DrpB, and DrpC. Of these, DrpC, which lacks many of the features required for Drp function, has been associated with mitochondrial division (14). Nonetheless, we and other groups have shown that instead DrpC appears to be involved in vesicle trafficking and endocytosis (14, 15). As the strongest homolog of any putative fission protein in *Toxoplasma* we investigated the role of Fis1 in monensin driven mitochondrial rearrangement. We found that Fis1 localization to the mitochondrion is important for monensin-induced remodeling and the absence of Fis1 results in decreased sensitivity to the ionophore. Thus, it is plausible that Fis1 is recruiting proteins to the mitochondrion outer membrane during monensin treatment to induce a transient constriction, similar to the transient interaction Fis1 has with Drp1 (34, 35). As DrpC and Fis1 do not seem to interact and DrpC localization does not change upon monensin treatment it is unlikely that DrpC is involved in this process. Interactome analysis of Fis1 identified some proteins with domains of interest that are also found in Fis1 interactors of other systems. For example, TGGT1_224270 contains WD40-like domains, which is common to the Fis1 adaptor proteins (34, 36). TGGT1_304990 is a guanylate-binding protein that may be able to take the role of a dynamin-related protein in this system.

While in yeast Fis1 is essential, mammalian cells appear to have several proteins able to recruit the fission machinery, which makes Fis1 dispensable in those organisms. Knockout of *Toxoplasma* Fis1 does not disrupt mitochondrial morphology (16) or affect parasite fitness (16, 28). These results are corroborated by our experiments in which the endogenous Fis1 gene was disrupted through CRISPR/Cas9 (Fig S1). Interestingly, we do observe a significant defect in the morphology of the mitochondrion when the endogenous Fis1 is mislocalized to the cytoplasm by deleting its transmembrane domain. In both mammalian cells and in yeast, either mislocalization or overexpression of Fis1 results in disruption of mitochondrial morphology (34, 37). In *Toxoplasma*, mislocalization of Fis1 resulted in aberrant mitochondrial morphology in which they maintain their lasso shape, but it is stretched out and appears to have strenuous branches and material. The phenotype observed with mislocalized Fis1 could the consequence of Fis1 interacting with proteins that it would normally not come into contact with or of Fis1 pulling proteins away from the mitochondrial membrane where they are required. With this in mind we performed a yeast two hybrid screen to identify putative interactors. Interestingly, among the 24 proteins identified, seven (TGGT1_215520, TGGT1_218560, TGGT1_265180, TGGT1_246720, TGGT1_304990, TGGT1_321370, and TGGT1_321450) likely localize to the mitochondrion, based on a proteomic analysis of the *Toxoplasma* mitochondrion, which uses both *BirA (38) and APEX (39, 40) to identify novel mitochondrial proteins (41). Nonetheless, this proteome may not contain all the potential interactors that localize to the mitochondrion because the proteome was generated using a mitochondrial matrix protein, HSP70, thus excluding proteins that are localized to the outer mitochondrial membrane. *In silico* analysis of the putative Fis1 interactors using MitoProt, SignalP, and PSort (42–44) shows that an additional 5 proteins (TGGT1_226050, TGGT1_237015, TGGT1_247700, TGGT1_299670, and TGGT1_286470) may also localize to the mitochondrion based on the presence of mitochondrial signal. Another protein of interest is TGGT1_287980 has a forkhead-associated (FHA) domain, which _i_s involved in a number of regulatory and signaling processes (45). Further characterization of these proteins is needed to determine what role they may play in mitochondrial remodeling and dynamics.

In this study, we focused on one of the putative Fis1 interactors, TGGT1_265180, which we have dubbed LMF1. This protein was the only to be identified through both the Y2H and a small-scale co-immunoprecipitation assay. LMF1 localizes to the OMM despite the absence of any domain or modification that would predict mitochondrial or membrane localization, suggesting that its association with the mitochondrion is likely through protein-protein interactions. When Fis1 is mislocalized to the cytoplasm, LMF1 expression is significantly reduced and while some LMF1 is still deposited on the mitochondrion, other remnants do not appear to be associated with the organelle. LMF1 may not colocalize with Fis1 in these parasites because either protein may be interacting with other proteins or membranes. In the case of LMF1, there are potentially redundant interactors on the mitochondrial surface or interactors localized to other parts of the parasite, like the IMC, that are important for maintaining the mitochondrial lasso shape. Additionally, the expression level of LMF1 is decreased significantly when Fis1 is mislocalized, which may be due to either a decrease in the transcript level of LMF1 or that the protein is being degraded in the absence of potentially stabilizing Fis1 interactions.

Genetic disruption of LMF1 reveals its unexpected role in maintenance of mitochondrial morphology in intracellular parasites. LMF1 knockout results in loss of the typical lasso arrangement with the majority of parasites having either sperm-like or collapsed mitochondria. Thus, it appears that in the absence of LMF1 the mitochondrion of intracellular parasites adopts morphology normally only seen in extracellular ones. These mitochondrial morphologies, sperm-like and collapsed, are proposed to be due to a retraction of the mitochondrion from the IMC as the parasite transitions to the extracellular environment. Therefore, it is possible that elimination of LMF1 has also eliminated these contact sites, causing a significant decrease in parasites with lasso morphology intracellularly. Membrane contact sites (MCSs) play important roles in signaling, lipid and ion exchange between organelles, and proper organelle positioning (46, 47). Whether any of these processes are affected in the LMF1 mutant strain is yet to be investigated. Nonetheless, the fact that parasites lacking LMF1 exhibit a propagation defect suggest that the proper morphology of the mitochondrion is important for parasite fitness.

We noted that complementation of the knockout strain with the wildtype LMF1 was incomplete. While the exogenous copy was under the control of the *LMF1* promoter, it is possible that the expression level from the ectopic site is not at the right level for complete complementation. Another possibility is that, in order to adapt to the lack of LMF1, the expression of other factors required for mitochondrial morphology was affected. Therefore, when the wildtype construct was introduced, it was missing other interactors that are necessary to fully attach the mitochondrion to the periphery. Future experiments using conditional knockout of LMF1 provide a better controlled system to study this mechanism.

Altering mitochondrial morphology is important in many systems to accommodate energetic needs and change positioning of organelles to perform specific functions. For example, mitochondria in lymphocytes concentrate towards the leading edge and alter their morphology to allow for chemotaxis to the site of injury (48). *Trypanosoma brucei* is another parasite that contains a single mitochondrion that alters its shape in different life stages (49). During the procyclic phase in the tsetse fly midgut, the mitochondrion elongates to form an elaborate network of mitochondrial branches. In the bloodstream form, the branches collapse to form one tubule that lacks the respiratory capability of the procyclic stage. This mitochondrial morphology change is dependent on a protein called TbLOK1, which is naturally downregulated in the bloodstream form (49). Based on this knowledge, it is possible that the retraction from the IMC toward the apical end of the parasite during extracellular stress is to a) position the mitochondrion to the area of greatest energetic need and/or b) accommodate to the available nutrients. We propose that LMF1 interacts with Fis1 on the OMM and another or multiple proteins in the parasite pellicle to establish membrane contact sites to maintain the typical lasso shape. Upon egress, LMF1 or its interactors is either post-translationally modified or downregulated as to eliminate these contact sites and position the mitochondrion towards the apical end. Once the parasite has reentered a host cell, the mitochondrion can then reattach to the pellicle and can extend to the parasite periphery. LMF1 knockout parasites cannot properly form this lasso and therefore have given us an incredible tool to study the functional relevance of the mitochondrial morphodynamics and to identify the key players in this process.

## EXPERIMENTAL PROCEDURES

### Host cell and parasite maintenance

All parasite strains were maintained via continued passage through human foreskin fibroblasts (HFF, purchased from ATCC) in normal growth medium, which consisted of Dulbecco’s modified Eagle’s medium (DMEM) supplemented with 10% fetal bovine serum (FBS), 2 mM L-glutamine, and 100 units penicillin/100µg streptomycin per mL. All cultures were grown in a humidified incubator at 37°C and 5% CO_2_. Parasites used were of the strain RH lacking hypoxanthine-xanthine-guanine phosphoribosyl transferase (HPT, RHΔ*hpt)* (50) and RH lacking HPT and Ku80 (RHΔ*ku80*Δ*hpt*, referred to as Δ*ku80* thereafter) (51, 52). For experiments involving drug treatment, the medium was supplemented with 1% FBS rather than 10%. For pyrimethamine treatment we used dialyzed serum. All drugs were purchased from Sigma. Stocks of monensin, pyrimethamine, and myxothiazol were prepared in ethanol, while atovaquone was prepared in DMSO.

### Generation of transgenic parasites

Parasites were engineered to express ectopic copies of full-length Fis1 (TGGT1_263323) or a truncated version lacking the putative transmembrane (TM) domain. For this purpose, PCR was utilized to amplify the Fis1 cDNA and append a hemagglutinin (HA)-tag at the N-terminus. The amplicon was flanked by NsiI and PacI restriction enzyme sites. Supplemental table S2 lists all the primers used throughout this study. Purified PCR fragments were inserted into the pHEX2 plasmid (53) using the In-Fusion HD Cloning Plus kit (Clontech). Expression of the transgenes was controlled by the *SAG1* promoter and selection was provided by the presence of the HPT selectable marker (50). 35µg of KpnI-linearized plasmids were electroporated into parental RHΔ*hpt* parasites (54) and selection of parasites that successfully integrated the plasmid was achieved by growing parasites in medium containing 50µg mycophenolic acid and 50µg xanthine per mL. Three rounds of drug selection were followed by limited dilution cloning to establish HA-tag positive parasite lines with and without the transmembrane domain termed RHΔ*hpt*+HAFis1 and RHΔ*hpt*+Fis1ΔTM, respectively.

To generate a parasite line expressing an endogenous Fis1 lacking the TM (RHΔ*ku80*:Fis1ΔTM), a fragment of the Fis1 gene comprising the region just upstream of the TM and flanked by PacI and AvrII was PCR amplified from *Toxoplasma* genomic DNA and inserted into the pLIC-HA(3x)-DHFR plasmid (51) by In-Fusion cloning. 35µg of EcoRV-linearized plasmid was transfected into Δ*ku80* parasites (51). Resulting transfectants were selected for dihydrofolate reductase (DHFR) by growth in medium with 1µM pyrimethamine and cloned by limited dilution.

For C-terminal endogenous epitope tagging of TGGT1_265180, a PacI-flanked fragment of TgGT1_265180 just upstream of its stop codon was PCR amplified and inserted into pLIC-myc(3x)-DHFR by In-Fusion cloning. 60µg of XcmI-linearized plasmid was transfected into Δ*ku80* parasites and transfectants were selected for DHFR as described above.

Double homologous replacement of the TGGT1_265180 coding sequence was used to establish a knockout strain. For this purpose, we generated a knockout construct using the previously described pminiGFP vector (55). Using In-Fusion cloning we introduced a 1,400bp PCR amplicon encompassing the region upstream of the TgGT1_265180 start codon into the HindIII restriction site of pminiGFP and a 1,156bp amplicon of the region downstream of the stop codon into the NotI restriction site. In this manner, the resulting vector (p265180_KO) has a drug selection cassette, HPT, flanked by regions of homology to the sequences upstream and downstream of TGGT1_265180. 10µg of DraIII-linearized p265180_KO was transfected into Δ*ku80* parasites using Nucleofector™ (Lonza) and parasites were then selected for the expression of HPT, as described above. Disruption of TGGT1_265180 was confirmed by PCR using three primer sets (Supplementary table S2). The first primer set (P1) amplifies a 637bp region present in wildtype parasites and absent in the knockout strain (Fig. 7A). The second primer set (P2) was designed to amplify a 1933bp fragment only present if the double homologous recombination of the knockout construct occurred at the TgGT1_265180 locus (Fig. 7A). The final primer set (P3) amplifies a fragment in both the wildtype and knockout strains (Fig. 7A).

For exogenous expression of TGGT1_265180, a 3700bp fragment beginning approximately 2kb upstream of the TgGT1_265180 start codon and ending at its stop codon was PCR amplified from genomic DNA. This PCR amplicon was inserted into the PacI site of pLIC-HA(3x)-DHFR by In-Fusion cloning. The same method was used to create a plasmid lacking the predicted SID, thus truncating the gene. These plasmids were used as templates to amplify an 8kb fragment that included the TgGT1_265180 gene under the control of its own promoter, a triple hemagglutinin tag, and the DHFR drug selection cassette. Primers used included overhangs homologous to the remnants of the Δ*ku80* site on each side of a double-stranded cut created by CRISPR/Cas9. The 8kb PCR fragment was gel extracted using the NucleoSpin Gel and PCR Clean-up kit (Macherey-Nagel) and eluted in P3 Buffer (Lonza) for nucleofection. The pSAG1-Cas9-U6-sgUPRT plasmid, generously provided by the Sibley lab (25), was mutated to contain a guide RNA targeted to the Ku80 site. TGGT1_265180 knockout and parental parasites were transfected with 1µg of either the full-length (265180-HA) or truncated (265180ΔSID-HA) PCR amplicons and 2µg of pSAG1-Cas9-sgKu80 using the Nucleofector™ (Lonza). Parasites were selected for the presence of DHFR, as described above. Immunofluorescence and western blot (see below) was used to confirm expression and localization of the exogenous copies of TgGT1_265180.

### Immunofluorescence microscopy analysis

For IFA, infected HFFs were fixed with 3.5% formaldehyde, quenched with 100 mM glycine, and blocked and permeabilized in 3% bovine serum albumin (BSA) and 0.2% Triton x-100 (TX-100) in PBS. Samples were then incubated with primary antibodies in PBS/3% BSA/0.2% TX-100 for one hour, washed five times with PBS, and incubated with Alexa Fluor conjugated secondary antibodies in PBS/3% BSA for one hour. Coverslips were washed with PBS and mounted on glass slides with 3 µL DAPI containing Vectashield. For 3D-SIM microscopy coverslips were stained with a liquid DAPI solution in PBS, washed, and inverted on a glass slide with Vectashield mounting medium without DAPI. Image acquisition and processing was performed on either a Nikon Eclipse 80i microscope with NIS-Elements AR 3.0 software or a Leica DMI6000 B microscope with LAS X 1.5.1.13187 software. 3D-SIM was performed utilizing the OMX 3D-SIM super-resolution system located within the Light Microscopy Imaging Center at Indiana University Bloomington (http://www.indiana.edu/~lmic/microscopes/OMX.html). The system is equipped with four Photometrics Cascade II EMCCD cameras that permit imaging four colors simultaneously and is controlled by DV-OMX software. Images processing was completed using the Applied Precision softWoRx software.

Primary antibodies used in this study included rabbit anti-HA (Cell signaling Technology), rabbit anti-myc (Cell Signaling Technology), a rabbit polyclonal antibody against the MORN1 protein (56), mouse monoclonal antibody 5F4 (detects F_1_B ATPase, P. Bradley, unpublished), and rabbit anti-acetyl-K40-α-tubulin (EMD Millipore ABT241), all used at 1:1,000, with the exception of 5F4 which was used at 1:5,000. Secondary antibodies included Alexa Fluor 594 or Alexa Fluor 488 conjugated goat anti-rabbit and goat anti-mouse (Invitrogen), all used at 1:2,000.

### Phenotypic characterization of mutant and complemented strains

For drug effects on mitochondrial morphology infected HFFs on coverslips were vehicle or drug treated with monensin (1 ng/mL), atovaquone (100 nM), pyrimethamine (1 µM), or myxothiazol (50 ng/mL) for 12 hours. To allow for recovery, drug medium was washed away and replaced with normal growth medium for an additional 12 hours. IFA was performed as above using F_1_B ATPase antibodies to monitor the mitochondrion. Samples were blinded and at least 100 vacuoles per sample were inspected. Experiments were performed in experimental and biological triplicates.

Plaque and doubling assays were performed with 12-well plates using standard methods (57). Briefly, for the plaque assays 500 freshly egressed parasites were added to confluent HFF monolaters. After four days of incubation, cultures were fixed with methanol for 5 minutes and stained with Crystal Violet. Plaques were imaged using a ProteinSimple imaging system and number of plaques were counted on a light microscope. Experiments were performed in experimental and biological triplicates.

### Yeast two-hybrid screen

Yeast two-hybrid screening was performed by Hybrigenics Services, S.A.S., Paris, France (http://www.hybrigenics-services.com). The coding sequence for Fis1 (aa 2-118; XM_018781322.1) was PCR-amplified and cloned into pB66 as a C-terminal fusion with the Gal4 DNA-binding domain (Gal4-Fis1). The construct was checked by sequencing and used as a bait to screen a random-primed *Toxoplasma* cDNA library constructed into pP6. pB66 derives from the original pAS2ΔΔ vector (58) and pP6 is based on the pGADGH plasmid (59). 46 million clones (5-fold the complexity of the library) were screened using a mating approach with YHGX13 (Y187 ade2-101::loxP-kanMX-loxP, mat*α*) and CG1945 (mat*α*) yeast strains as previously described (58). 247 His+ colonies were selected on a medium lacking tryptophan, leucine and histidine. The prey fragments of the positive clones were amplified by PCR and sequenced at their 5’ and 3’ junctions. The resulting sequences were used to identify the corresponding interacting proteins in the GenBank database (NCBI) using a fully automated procedure. A confidence score (PBS, for Predicted Biological Score) was attributed to each interaction as previously described (60).

### Immunoprecipitation Assay

To confirm the results of the yeast two-hybrid screening, we performed one immunoprecipitation assay using RHΔ*hpt*+HAFis1, with the parental RHΔ*hpt* parasites as a negative control. Extracellular parasites from 10 T175 culture flasks were spun down, washed twice with cold PBS, and resuspended in Pierce Co-IP Lysis buffer (Fisher Scientific) with Protease/Phosphatase Inhibitor Cocktail (100X, Cell Signaling Technology). After one hour of lysis at 4°C, the samples were sonicated three times for 15 seconds, with one-minute rest period between each sonication. After sonication, samples were pelleted and the supernatant transferred to Pierce™ Anti-HA Magnetic Beads (Fisher Scientific). Samples were placed on a rocker at 4°C for 2.5 hours before beads were washed once with Pierce Co-IP Lysis buffer and twice with PBS. Beads were resuspended in 8M urea and sent for LC/MS-MS analysis. Results were narrowed down to proteins that had at least 4 peptides in the RHΔ*hpt*+HAFis1 sample and none in the RHΔ*hpt* control. This shortened list was then compared to the list of putative interactors obtained through yeast two-hybrid.

### Western blots

Extracellular parasites were pelleted and resuspended in 2X Laemmli Sample Buffer (Bio-Rad) with 5% 2-mercaptoethanol (Sigma-Aldrich). Samples were boiled for 5 minutes at 95°C before separation on a gradient 4-20% SDS-PAGE gel (Bio-Rad). Samples were then transferred to nitrocellulose membrane using standard methods for semi-dry transfer (Bio-Rad). Membranes were probed with rabbit anti-HA (Cell Signaling Technologies), mouse anti-c-myc (Cell Signaling Technologies), or mouse anti-SAG1 (Thermo Fisher) at a dilution of 1:5000 for 1 hour. Membranes were then washed and probed with either goat anti-mouse horseradish peroxidase or goat anti-rabbit horseradish peroxidase (Sigma-Aldrich) at a dilution of 1:10000 for 1 hour (GE Healthcare). Proteins were detected using SuperSignal West Femto substrate (Thermo Fisher) and imaged using the FluorChem R system (Biotechne). All original western blots are shown in supplemental dataset 2.

For comparative analysis of LMF1 protein levels in RHΔ*ku80*:Fis1ΔTM parasites to that of RHΔ*ku80*, parasites were centrifuged and washed once with PBS. Parasites were counted using a hemocytometer and the parasite pellets were resuspended at appropriate volumes to equilibrate the concentration of parasites. The subsequent immunoblots were then probed for anti-SAG1 as a loading control. ImageJ was used for densitometry analysis of the detected protein band and compared to SAG1 signal. The ratio of LMF1 protein levels (normalized to the SAG1 levels in the same sample) of RHΔ*ku80*:Fis1ΔTM to RHΔ*ku80* was determined and represented as a percentage. These were done in biological triplicate and the described percentage is an average of these replicates.

### Statistical analysis

Statistics were performed with either JMP14.0 or Prism software.

## ACKNOWLEDGMENTS

This work was by grants from the National Institutes of Health to GA (R01AI123457 and R21AI138255). KJ is funded by a fellowship from NRSA training grant T32AI060519. RC was funded by a fellowship from the American Heart Association (15POST22740002). The funders had no role in study design, data collection and interpretation, or the decision to submit the work for publication. We would like to thank Hybrigenics for their analyses and support using Y2H. We would also like to thank Dr. Peter Bradley for generously providing F_1_B ATPase antibody.

